# A three-dimensional human adipocyte model of fatty acid-induced obesity

**DOI:** 10.1101/2022.02.24.481624

**Authors:** Vera M. Pieters, Saifedine T. Rjaibi, Kanwaldeep Singh, Nancy T. Li, Safwat T. Khan, Sara S. Nunes, Arianna Dal Cin, Penney M. Gilbert, Alison P. McGuigan

## Abstract

Obesity prevalence has reached pandemic proportions, leaving individuals at high risk for the development of diseases such as cancer and type 2 diabetes. In obesity, to accommodate excess lipid storage, adipocytes become hypertrophic, which is associated with an increased pro-inflammatory cytokine secretion and dysfunction of metabolic processes such as insulin signaling and lipolysis. Targeting adipocyte dysfunction is an important strategy to prevent the development of obesity-associated disease. However, it is unclear how accurately animal models reflect human biology, and the long-term culture of human hypertrophic adipocytes in an *in vitro* 2D monolayer is challenging due to the buoyant nature of adipocytes. Here we describe the development of a human 3D *in vitro* disease model that recapitulates hallmarks of obese adipocyte dysfunction. First, human primary adipose-derived stromal cells are embedded in hydrogel, and infiltrated into a thin cellulose scaffold. The thin microtissue profile allows for efficient assembly and image-based analysis. After adipocyte differentiation, the scaffold is stimulated with oleic or palmitic acid to mimic caloric overload. Using functional assays, we demonstrated that this treatment induced important obese adipocyte characteristics such as a larger lipid droplet size, increased basal lipolysis, insulin resistance and activation of macrophages through adipocyte-conditioned media. This 3D disease model mimics physiologically relevant hallmarks of obese adipocytes, to enable investigations into the mechanisms by which dysfunctional adipocytes contribute to disease.

## Introduction

Obesity is a medical condition, defined by a body mass index of 30 kg/m^2^ or over and associated with an increased risk of development of various comorbidities such as osteoarthritis, cancer, cardiovascular disease and type 2 diabetes.^1^ It has been well established that obesity prevalence is on the rise and it is estimated that 42 % of the world’s population will live with obesity by 2030.^2^ Obesity is characterized by an expansion of white adipose tissue. White adipose tissue is composed of adipocytes, the lipid-storing cells, and a large stromal vascular fraction consisting of precursor cells, fibroblasts, endothelial cells and an immune cell population.^3^ In obesity, due to their role in lipid storage, adipocytes are one of the first cell types affected by a positive energy balance,^4^ which highlights the importance of investigating the role of obese adipocytes in the development of metabolic disorder.

In obesity, adipocytes store excess lipids in lipid droplets in the form of triglycerides, leading to significant increase in lipid droplet and cell size.^5^ Lipid droplet hypertrophy is associated with insulin resistance, which increases the levels of glucose and triglycerides in the circulation.^6^ Additionally, hypertrophic adipocytes express increased rates of lipolysis, a process in which triglycerides are released from the cells in the form of free fatty acids and glycerol, in the absence of catecholamine stimulation.^7^ In healthy adipocytes, these metabolic processes of energy uptake and release are tightly regulated, but in obesity, impairment of these mechanisms leads to metabolic disorder.^8^ Moreover, while adipocytes were previously thought of as mere lipid storages, over the recent years they have been recognized for their important secretory function. Specifically, hypertrophic adipocytes secrete higher levels of pro-inflammatory cytokines, which contributes to chronic, low-grade inflammation on a local and systemic level.^9,10^

It is difficult to study disease mechanisms of obese adipocytes *in vivo* due to the multicellular complexity of adipose tissue and the challenge of correlating specific microenvironmental factors to the cellular phenotype. Conversely, simpler 2D cell culture strategies are unsuitable for long term adipocyte differentiation and culture: adipocytes are large, spherical and contain a high volume of lipids which makes them buoyant and leads to cell detachment from the cell culture plate.^11^ Additionally, tissue culture plastic is not ideal for adipocyte culture because adipocytes express a fibrotic phenotype and a decreased adipogenic potential when in contact with a stiff plastic surface.^12,13^ To investigate the mechanisms by which obese adipocytes contribute to development of metabolic disorder in people with obesity, we set out to create an obese adipocyte model that overcomes the aforementioned challenges.

We expected it would be essential to establish a protocol that mimics a positive energy balance to induce the characteristics of obese adipocytes to investigate the mechanisms by which obese adipocytes contribute to metabolic disorder. Consistent with our hypothesis, previous studies demonstrated that stimulation of 2D monolayers of murine adipocytes with fatty acids induces obesity characteristics such as lipid droplet hypertrophy, insulin resistance and an altered cytokine profile.^14,15^ Other groups have stimulated multicellular self-assembled spheroids with a lipid mix and demonstrated similar obesity characteristics.^11,16^ However, 2D strategies do not mimic the complex 3D architecture that the cells are exposed to *in vivo*, while multicellular self-assembled spheroids do not enable control over the density and size of the adipose microtissue to understand how organization influences tissue function. Further, while it was shown in murine adipocytes that saturated fatty acids induce a different phenotype in adipocytes (increased pro-inflammatory cytokine production, disruption of circadian rhythm, increased endoplasmic reticulum stress) as compared to unsaturated fatty acids,^14,17–19^ it remains unclear what the effect is of saturated and unsaturated fatty acids on inducing obesity characteristics in human adipocytes.

As such, we describe a strategy to culture human adipocytes in 3D that enables controlled assembly and easy imaging and in which we can induce an obese phenotype by mimicking caloric overload via stimulating with different fatty acids. By adapting a culture platform previously developed by our group^20–27^, we infiltrated human adipose-derived stromal cells (ADSCs) suspended in a hydrogel into a porous cellulose scaffold and differentiated the cells into mature adipocytes. This resulted in a thin, but three-dimensional adipocyte culture system that has the benefit of providing the cells with a 3D environment, suitable for long term adipocyte culture, while still enabling easy image-based analysis and simple modification of microenvironmental factors. In order to mimic caloric overload, we stimulated the scaffolds with either oleic acid, an unsaturated fatty acid, or palmitic acid, a saturated fatty acid, over a period of up to a week. Using functional assays, we assessed adipocyte lipid droplet hypertrophy, insulin resistance, basal lipolytic rate and activation of macrophages through adipocyte-conditioned media. We found that these obesity characteristics were increased after stimulation with both fatty acids, with palmitic acid inducing higher rates of lipolysis and a significant change in macrophage activation. Our approach provides a new strategy to mimic caloric overload in a human 3D adipocyte culture system which will support studies to elucidate the mechanisms by which obese adipocytes contribute to metabolic disorder.

## Methods

### Human ethics

The human adipose-derived stromal cells used for the experiments demonstrated in **Figure 1b, 1c, 2, 3, 4, and 5, and Supplementary Information 1, 3b, 5, 7, 8, 9, and 10** were derived from adipose tissue collected during an elective bilateral mastectomy conducted by Dr. Arianna Dal Cin under McMaster University Research Ethics Board approval (REB#13078-T). Only adipose tissue removed during scheduled surgical procedures and designated for disposal were utilized in this study. Informed consent was obtained from all study participants. The University of Toronto Office of Research Ethics reviewed the approved studies and further assigned administrative approval (Protocol# 00042225). All methods in this study were performed in accordance with the guidelines and regulations of these two Research Ethics Boards.

**Figure 1.**
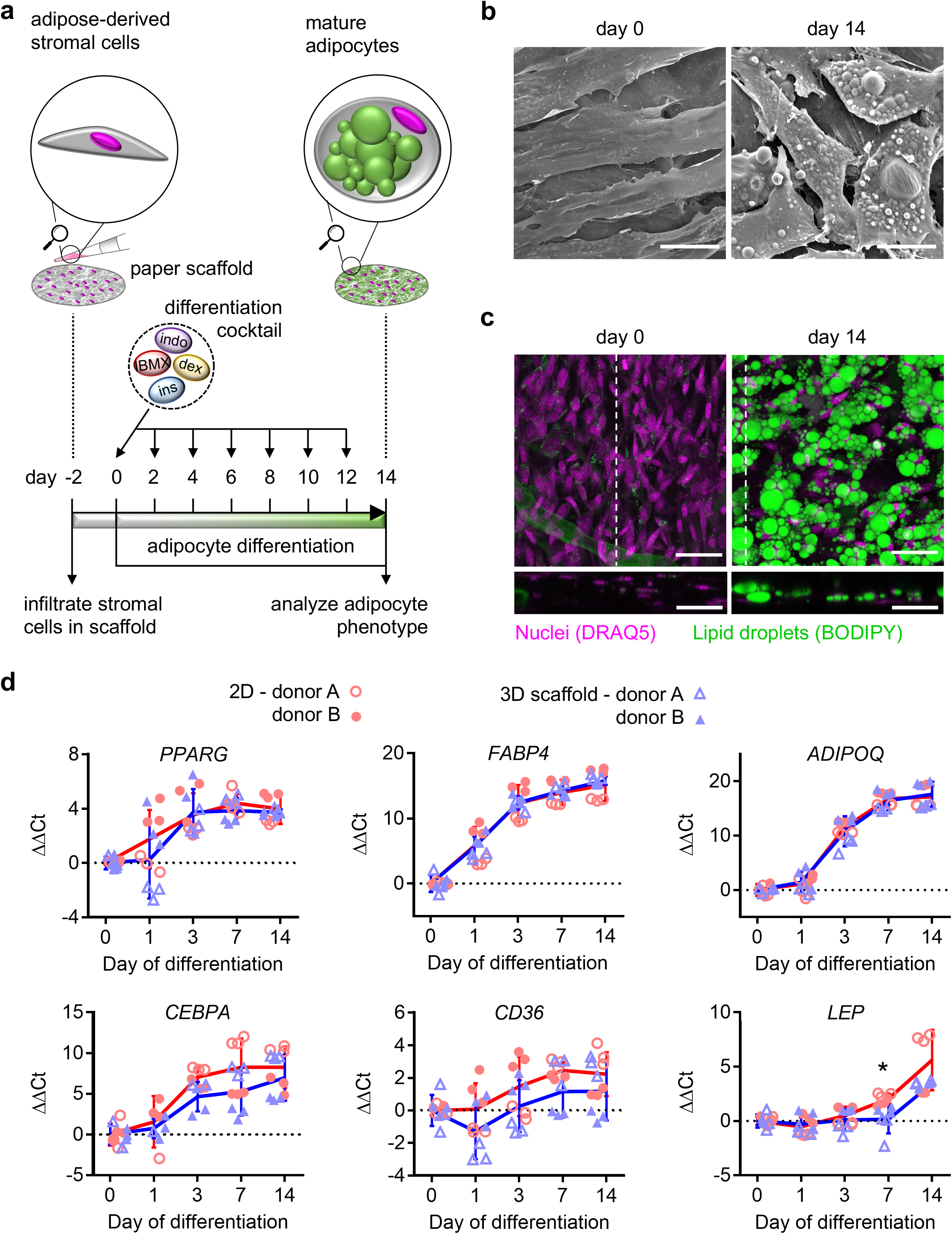
Differentiation of human adipose-derived stromal cells into adipocytes in a 3D paper scaffold. **a.** Experimental timeline of adipocyte differentiation in 3D paper scaffold. **b.** Representative scanning electron microscopy images of 3D paper scaffold with human adipose-derived stromal cells at day 0 and day 14 of differentiation. Magnification 750x; scale bar, 25 μm. **c.** Representative confocal images and the respective side profiles of 3D paper scaffolds containing human adipose-derived stromal cells at day 0 and day 14 of differentiation. White dotted line displays the location of the side profile in the sample. Scaffolds were stained with the lipid-soluble dye BODIPY (green) and DNA stain DRAQ5 (magenta). Magnification 20x; scale bar, 50 μm. **d.** RT-qPCR analysis of adipocyte associated genes showing the comparison adipocyte differentiation in 2D (red circles, averaged by red line) and in 3D paper scaffold (blue triangles, averaged by blue line) culture. Data is displayed as ΔΔCt relative to day 0 of differentiation, using the geometric mean of reference genes *RPL13A* and *TBP.* Experiments were performed with cells derived from two different human donors, identifiable by the open or closed data points. Each data point represents the average of three RT-qPCR technical replicates of either one 2D well or one 3D paper scaffold. Error bars represent standard deviation. n=6 (3 wells/time point/cell donor for 2D culture and 3 scaffolds/time point/donor for 3D paper scaffold culture). Statistics performed by two sample T tests with Bonferroni correction. *p <0.01.

**Figure 2.**
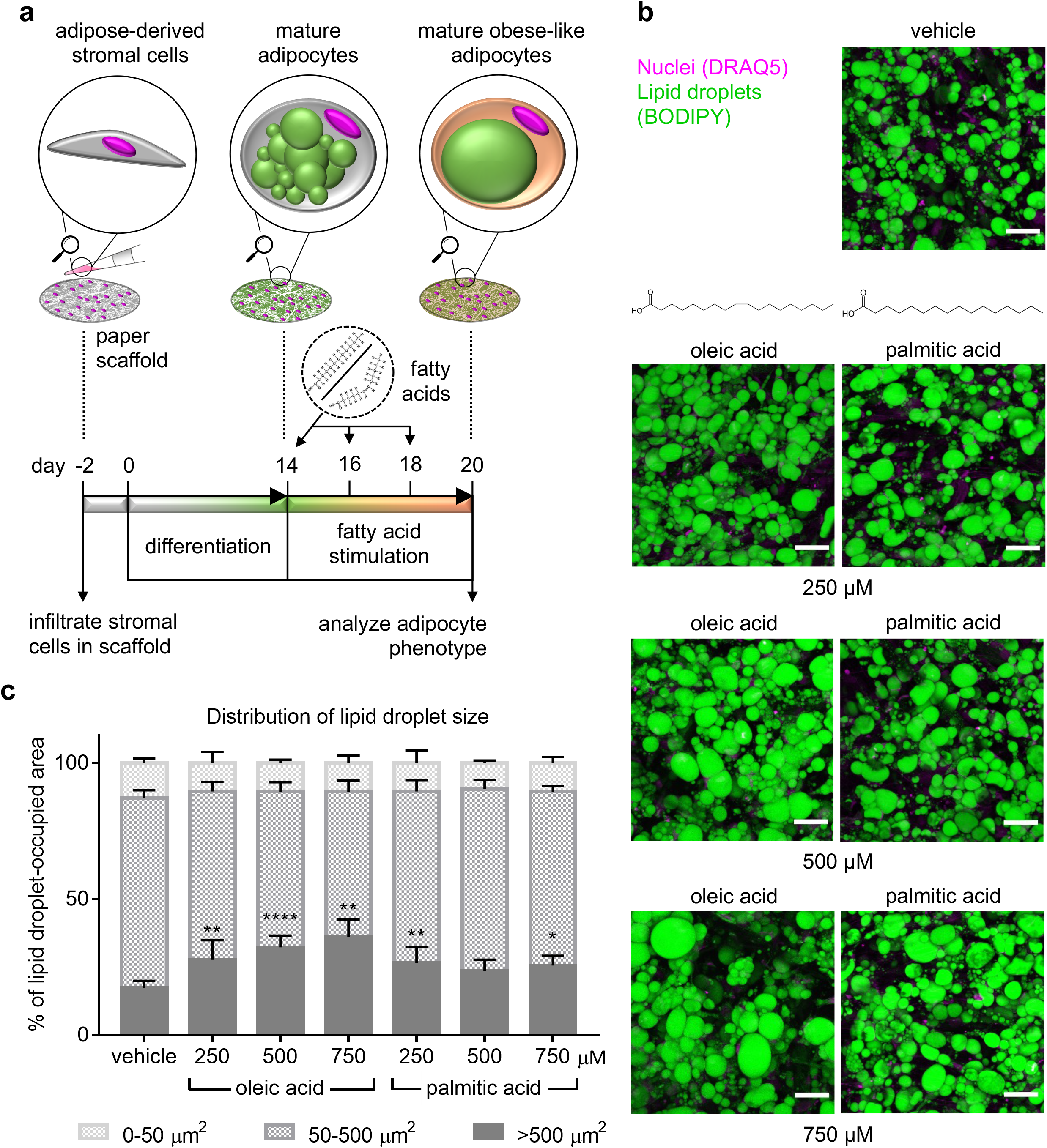
Fatty acid stimulation of human adipocytes in 3D paper scaffold increases lipid droplet size. **a.** Experimental timeline of fatty acid stimulation of human adipocytes in 3D paper scaffold. **b.** Representative confocal images of 3D paper scaffolds containing human adipocytes stimulated with oleic acid or palmitic acid in various concentrations for 6 days. Scaffolds were stained with the lipid-soluble dye BODIPY (green) and DNA stain DRAQ5 (magenta). Magnification 20x; scale bar, 50 μm. **c.** Quantification of lipid droplet size in human adipocytes in 3D paper scaffold stimulated with oleic acid or palmitic acid in various concentrations for 6 days. Lipid droplets were grouped in different size categories of 0-50 μm^2^ (light grey), 50-500 μm^2^ (medium grey), and >500 μm^2^ (dark grey). Data is displayed as the percentage of lipid droplet area that is occupied by lipid droplets in the three different size categories. Experiments were performed with cells derived from two different human donors. Each data point represents the average of two images of a 3D paper scaffold. Error bars represent standard deviation. n=6 to 9 (2 to 3 scaffolds/stimulation condition/donor). Statistics performed by one-way ANOVA and post-hoc Dunnet’s test, comparing the lipid droplet-occupied area in the >500 μm^2^ size category of each fatty acid stimulated group to the vehicle control. *p <0.05; **p<0.01; ****p<0.0001.

**Figure 3.**
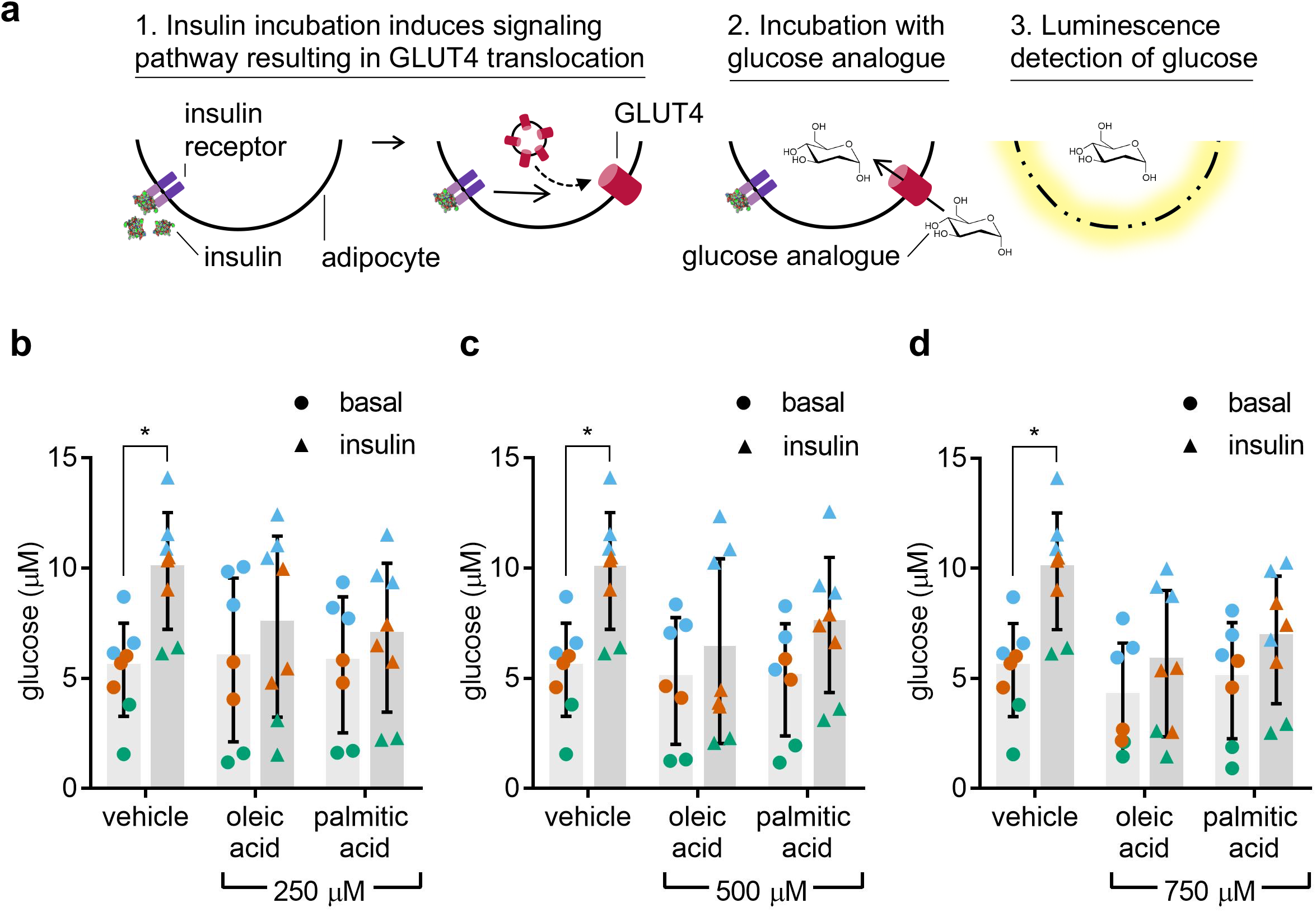
Fatty acid stimulation of human adipocytes in 3D paper scaffold impairs insulin sensitivity. **a.** Schematic of cellular mechanism of glucose uptake by the 3D adipocyte paper scaffold in response to insulin, assessed by a functional insulin sensitivity assay. After incubation with insulin, insulin-sensitive adipocytes initiate a signaling pathway that leads to fusion of GLUT4-loaded vesicles with the plasma membrane. The adipocytes are then incubated with a glucose analogue that is transported into the cell by GLUT4 transporters. At the assay endpoint, after a series of chemical reactions, the adipocytes are lysed and the released glucose analogue gives rise to a luminescent signal that is measured in a plate reader. **b-d.** Luminescence assessment of basal and insulin-stimulated glucose uptake of 3D adipocyte scaffolds after stimulation with **(b)** 250 μM, **(c)** 500μM, or **(d)** 750μM of oleic or palmitic acid. 3D adipocyte scaffolds were incubated in either the presence (triangles) or absence (circles) of 10 μM human insulin. Data is displayed as concentration of glucose taken up by the cells in each scaffold, normalized by a fluorescent measurement of cell viability of each scaffold. Three experiments were performed with cells from one human donor, identifiable by the green, orange or blue data points. Each data point represents a 3D adipocyte scaffold. Statistics performed by two-way ANOVA and post-hoc Sidak’s test. *p <0.05.

**Figure 4.**
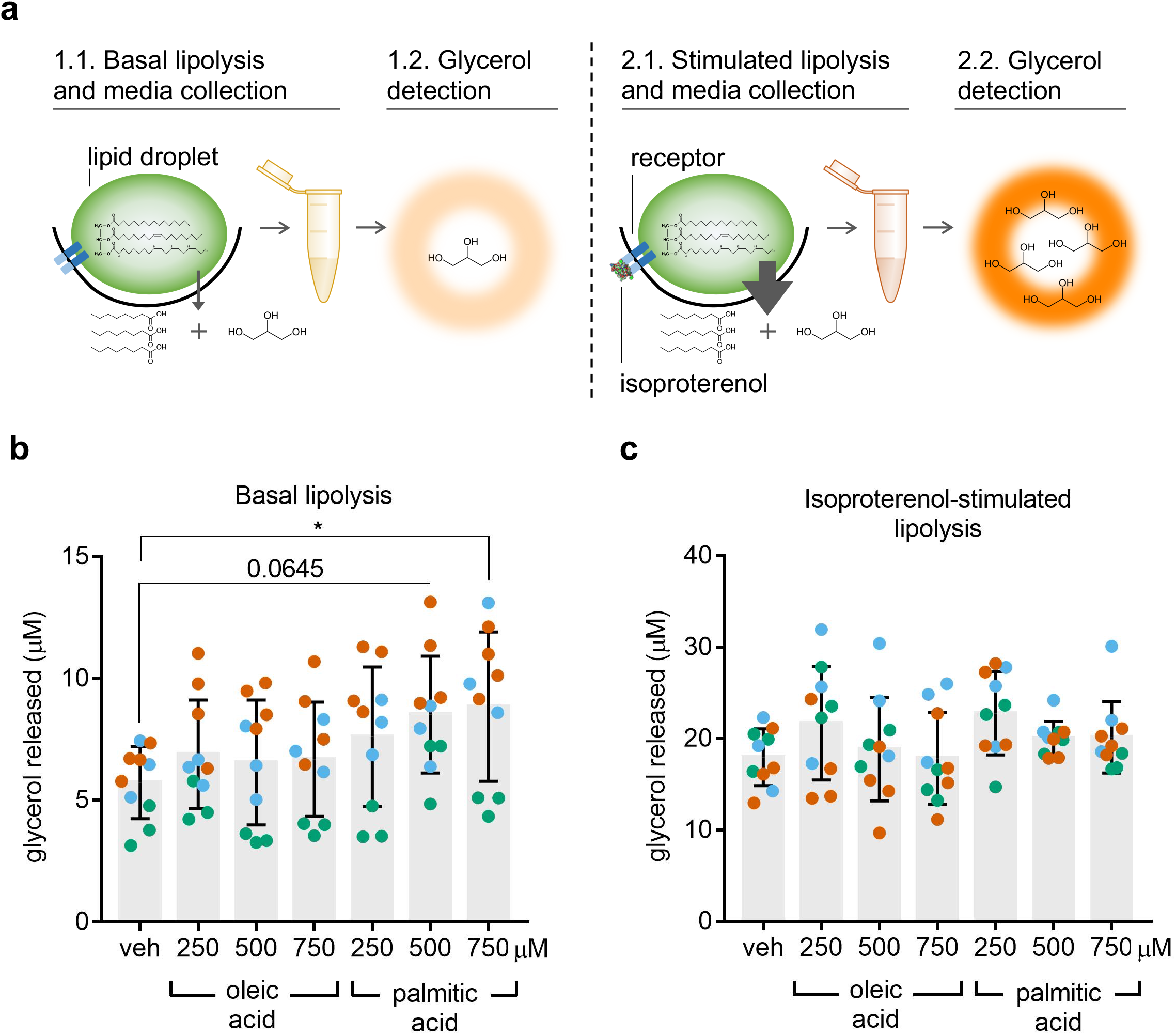
Palmitic acid stimulation of human adipocytes in 3D paper scaffold increases lipolysis. **a.**Schematic of cellular mechanism of lipolysis in the 3D adipocyte paper scaffold. Lipolysis is the secretion of triglycerides from the lipid droplets into free fatty acids and glycerol. This assay collects conditioned media from adipocytes before (basal) and after β3 adrenergic stimulation. As a result of a series of chemical reactions, the glycerol released in the media gives rise to a luminescent signal that is measured in a plate reader, as a measurement for lipolysis. **b.** Luminescence assessment of basal glycerol release of 3D adipocyte scaffolds after stimulation with various concentrations of oleic or palmitic acid. Data is displayed as concentration of glycerol released by the cells in each scaffold, normalized by a fluorescent measurement of cell viability of each scaffold. Three experiments were performed with cells from one human donor, identifiable by the green, orange and blue data points. Each data point represents a 3D adipocyte scaffold. **c.** Luminescence assessment of isoproterenol-incubated glycerol release of 3D adipocyte scaffolds after stimulation with various concentrations of oleic or palmitic acid. Data is displayed as concentration of glycerol released by the cells in each scaffold, normalized by a fluorescent measurement of cell viability of each scaffold. Three experiments were performed with cells from one human donor, identifiable by the green, orange or blue data points. Each data point represents a 3D adipocyte scaffold. Statistics performed by oneway ANOVA and post-hoc Dunnet’s test. *p <0.05.

**Figure 5.**
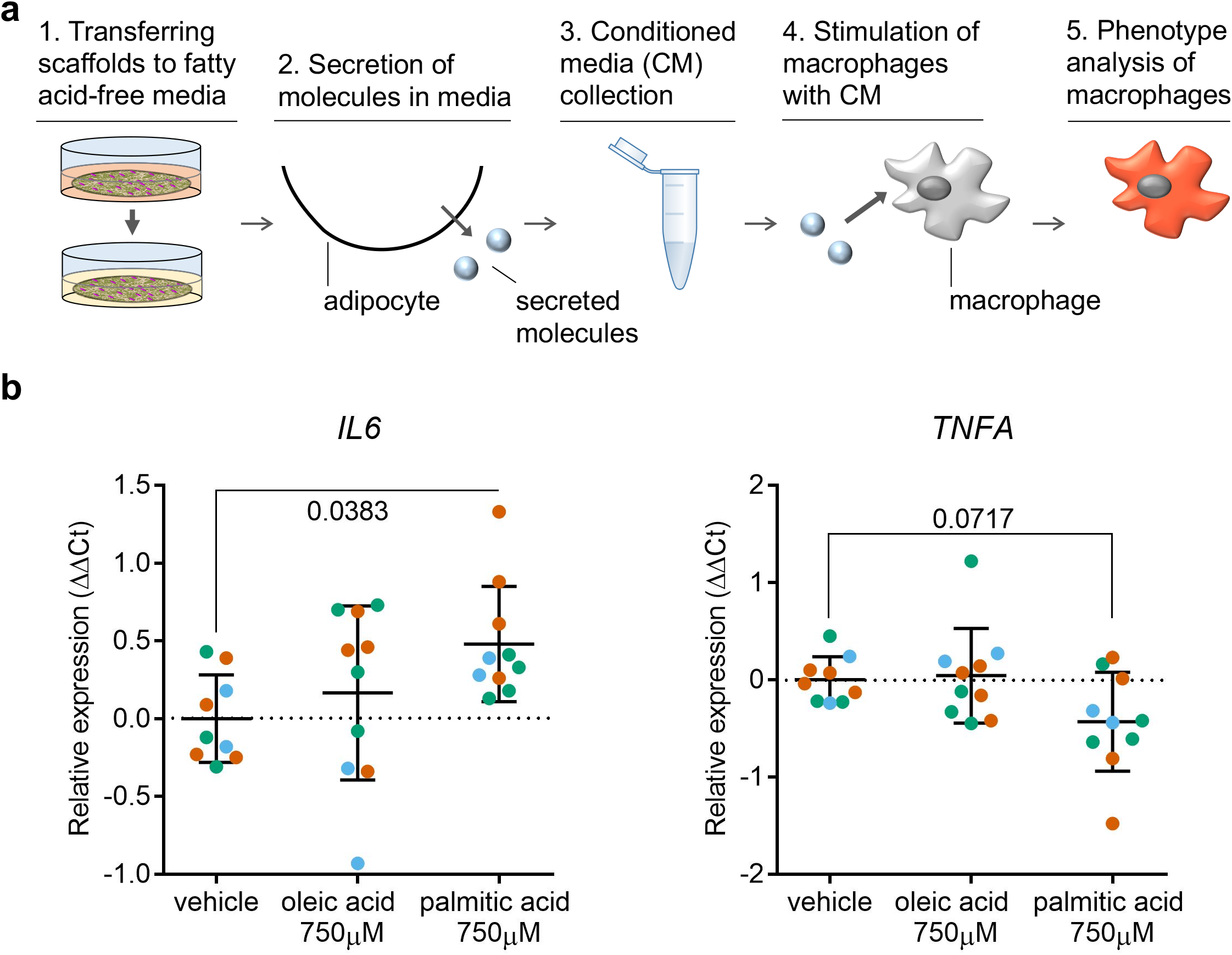
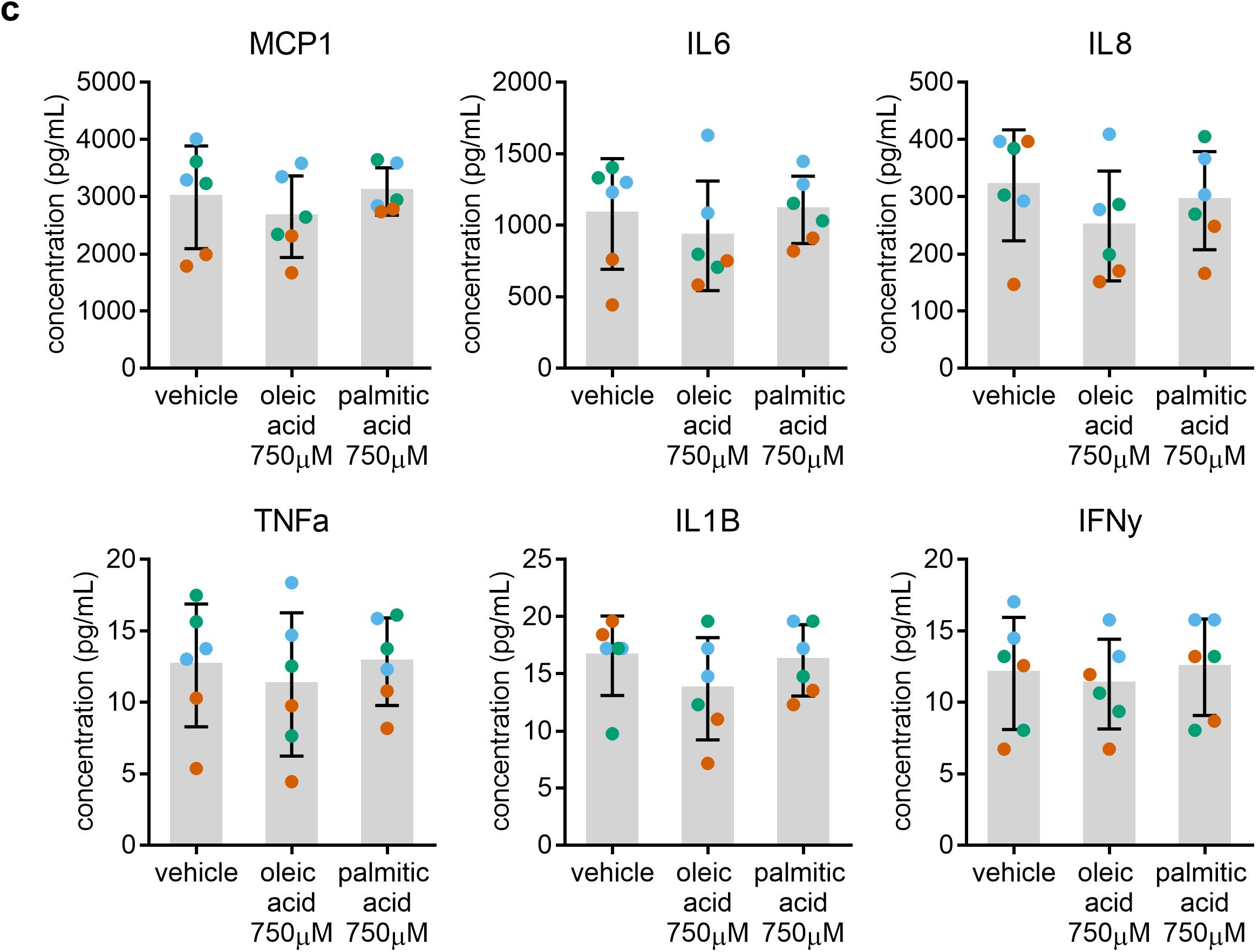
Fatty acid stimulation of human adipocytes in 3D paper scaffold does not affect their pro-inflammatory cytokine profile, but does alter macrophage phenotype through adipocyte-conditioned media. **a.** Schematic of experimental procedures of stimulating macrophages with fatty acid-stimulated 3D adipocyte scaffold-conditioned media. Primary human macrophages are stimulated with adipocyte-conditioned media. The effect of the adipocyte-conditioned media on macrophages is assessed by analyzing gene expression of pro-inflammatory macrophage markers with qPCR. **b.** RT-qPCR analysis of pro-inflammatory gene expression in macrophages in response to stimulation with adipocyte-conditioned media. Three experiments were performed with adipocyte-conditioned media from one human donor, identifiable by the green, orange or blue data points. Data is displayed as ΔΔCt relative to vehicle, using the geometric mean of reference genes *RPL13A* and *TBP*. Experiments were performed with cells derived from one human macrophage donor. Each data point represents the average of three RT-qPCR technical replicates of one 2D well of macrophages, stimulated with conditioned media from one 3D paper scaffold. Error bars represent standard deviation. n=9 (for vehicle) or 10 (for fatty acid-stimulated adipocyte conditioned media). Statistics performed by one-way ANOVA and post-hoc Dunnet’s test. P values are described in the figure. **c**. Multiplex analysis of cytokines released into the cell culture media from 3D adipocyte scaffolds in response to stimulation with fatty acids. Three experiments were performed with adipocytes derived from one human donor, identifiable by the green, orange and blue data points. Data is displayed as the concentration of released cytokines into the cell culture media (pg/mL). Each data point represents the average of two measurements of the cell culture media from one 3D adipocyte scaffold. Error bars represent standard deviation. n= 6. Statistics performed by one-way ANOVA and post-hoc Dunnet’s test, no significance p values were observed.

### Adipose-derived stromal cell derivation and maintenance

The P21.5 primary human adipose-derived stromal cell (hADSCs) line we used in this study was derived from adipose biopsy tissue (P21.5). Briefly, adipose tissue was minced into 1-2 mm pieces and the minced tissue was digested with 200 U/mL Collagenase I (ThermoFisher) at 37 °C in a shaking water-bath for 1 hour, and the degree of digestion was checked every 10 minutes. The digested tissue was filtered using a 100 μm nylon filter to remove undigested tissue from the liberated cells. The filtrate was centrifuged at 300 *g* for 10 minutes to pellet stromal vascular fraction containing the hADSCs and cultured in ADSC growth media for 2 weeks with media change every 3 – 4 days. hADSCs were checked for fibroblast like morphology, adherence to regular cell culture plasticware and expression of mesenchymal stem cell markers, CD73 (BD Biosciences, Cat# 561254), CD90 (BD Biosciences, Cat# 555596) and CD105 (BD Biosciences, Cat# 562408) by flow cytometry. The primary subcutaneous adipose-derived stromal cell lines designated herein as Z25.3, Z24.5 were purchased from a commercial source (ZenBio). Donor information is described in **Supplementary Information Table 1**. The donors had no history of diabetes or cancer and did not take any medications. hADSCs were plated in 75 cm^2^ tissue culture treated flasks at a density of ~5000 cells/cm^2^ in growth media consisting of DMEM/F12 with L-Glutamine and 2.493 g/L sodium biocarbonate (Gibco) supplemented with 10 % fetal bovine serum (Wisent Bioproducts) and recombinant human FGF-basic at a concentration of 10 ng/mL (Peprotech). hADSCs were passaged upon reaching ~80 % confluency, using Trypsin-EDTA (Wisent Bioproducts). hADSCs between passage 5 and 8 were used for adipocyte scaffold creation or adipocyte differentiation in 2D. For adipocyte differentiation in 2D, hADSCs were plated onto 48-well plates in growth media. When confluent, growth media was aspirated and replaced by adipocyte differentiation media consisting of DMEM/F12, supplemented with 10 % FBS, 250 μM 3-isobutyl-1-methylxanthine (Sigma-Aldrich), 100 μM indomethacin (Sigma-Aldrich), 0.5 μM dexamethasone (Sigma-Aldrich) and 10 μg/mL human insulin (Sigma-Aldrich). Adipocyte differentiation media was replaced every two days until day 14.

### Adipocyte scaffold creation

Scaffold discs were created by punching discs with a 5.0 mm diameter biopsy punch (Integra Miltex) out of a commercially available cellulose scaffold product (MiniMinit^®^ Products Ltd., One Cup Filter). The cellulose discs were sterilized by autoclaving and placed onto a PDMS slab using forceps. The PDMS slab was then placed into humidification chamber, consisting of a plastic container with a paper towel on the bottom, humidified by adding 10 mL sterile PBS and closed with a lid.

A solution of fibrinogen (Sigma-Aldrich) was prepared in a NaCl solution (0.9 %wt/vol in ddH_2_O) at a concentration of 10 mg/mL and sterilized by passing it through a 0.22 μm filter. The extracellular matrix (ECM) master mix was created by combining 40 % DMEM/F12, 40 % 10 mg/mL fibrinogen and 20 % Geltrex™ (Gibco) and kept on ice. hADSCs were suspended in the ECM master mix at a concentration of 19 × 10^6^ cells/mL and kept on ice. 100 U/mL of thrombin (Sigma) was added to the hADSC-ECM mixture to produce a final concentration of 0.225 U/mg of fibrinogen. All cell culture solutions used in this work are summarized in **Supplementary Information Table 2**.

The humidification chamber was opened and 5 μL of the hADSC-ECM mixture (containing 90,000 cells) was pipetted onto the cellulose scaffold disc and infiltrated in between the fibers of the scaffold via capillary wicking. The humidification chamber was closed by returning the lid and the chamber was placed into an incubator set at 37°C for 5 minutes during which the fibrinogen converts to fibrin clots. Sterile PBS was pipetted onto the scaffolds to enable detaching the scaffolds from the PDMS slab using forceps. The scaffolds were transferred to individual wells of a 96-well tissue culture plate containing hADSC growth media. After 2 days, growth media was aspirated and replaced by adipocyte differentiation media consisting of DMEM/F12, supplemented with 10 % FBS, 250 μM 3-isobutyl-1-methylxanthine, 100 μM indomethacin, 0.5 μM dexamethasone and 10 μg/mL human insulin. Adipocyte differentiation media was replaced every two days until day 14.

### Stimulation of adipocyte scaffolds with fatty acids

We purchased palmitic acid (Cayman Chemicals) and oleic acid (Cayman Chemicals) solutions that were pre-conjugated to BSA at a ratio of 6:1. This fatty acid/albumin ratio mimics pathophysiological conditions.^28^ Fatty acid media was made in DMEM/F12 supplemented with 1 % FBS in concentrations of 250 μM, 500 μM or 750 μM of fatty acids and passed through a 0.22 μm filter to ensure sterility. Vehicle media was made by dissolving 80 μM of fatty acid-free BSA (Sigma-Aldrich) in DMEM/F12 supplemented with 1 % FBS. At day 14 of differentiation, scaffolds were washed with PBS and incubated with fatty acid media or vehicle media. Media was replaced every 2 days for a total of 6 days of fatty acid stimulation, with the exception of **Supplementary Information Figure 3** where we used 4 and 8 days of fatty acid stimulation.

### Scanning electron microscopy

Scaffolds were washed with PBS and fixed with 4 % paraformaldehyde (Thermo Scientific) and 2 % glutaraldehyde (Sigma-Aldrich) in PBS for 30 minutes at room temperature and then transferred to a solution of 1 % paraformaldehyde and 0.5 % glutaraldehyde overnight at room temperature. The scaffolds that contained cells were incubated in 1 % osmium tetroxide in PBS for 15 minutes at room temperature. All scaffolds were washed twice with PBS and placed in a cryovial. The cryovial was placed in liquid nitrogen for 1 minute to snap-freeze the samples. A vial containing 100 μL of PBS was included as a blank control to monitor the snap-freezing and freeze-drying processes. The cryovials were each opened slightly and placed in a freeze-dryer (Flexi-Dry MP, FTS Systems) for 2 hours. Samples were mounted onto carbon tape-coated stubs and gold–palladium sputter coated for 55 seconds using a Bal-Tec SCD050 Sample Sputter Coater (Leica Biosystems, USA). Images were obtained at 10 kVD using a Hitachi SEM SU3500 (Hitachi High-Technologies Canada Inc., Toronto, Canada).

### Lipid droplet imaging

Scaffold discs were washed with PBS and fixed with 4 % paraformaldehyde in PBS for 10 minutes at room temperature. Paraformaldehyde was aspirated and the scaffolds were incubated in the staining solution consisting of 1 μg/mL BODIPY (Invitrogen) and 5 μM DRAQ5 (Cell Signalling Technology) in PBS for 30 minutes at room temperature. The scaffolds were washed twice with PBS, transferred to a microscopic slide, mounted with Fluoromount-G (SouthernBiotech) and covered with a cover glass. Samples were imaged within 24 hours with an SP8 Confocal Microscope (Leica), equipped with a 20x objective (NA 0.75). Excitation and emission wavelengths were set to 503/512 nm for BODIPY fluorescence and 647/683 nm for DRAQ5 fluorescence. Each confocal image consisted of 35 optical slices with a dimension of 584 x 584 μm. Two to three images were taken of each scaffold.

### Lipid droplet size quantification

Lipid droplets were quantified using a custom automated image analysis workflow developed using the FIJI software. First, 3D confocal stacks were transformed to the 2D space using a Maximum Intensity Projection. The projected image was sharpened using a 3×3 filter kernel with center element “15” and all other kernel elements “-1”. Next, FIJI’s Moments thresholding algorithm was applied to binarize the filtered image. The resulting objects were segmented to obtain individual lipid droplets using FIJI’s Watershed tool, followed by three iterations of Openings (count=7) to clean up any background pixels, jagged edges, or dark pixels within the lipid droplets as a result of upstream processing. The finalised mask was fed to FIJI’s Analyze Particles tool where the area of each lipid droplet and total number of lipid droplets in the image were extracted. Objects that were cut off by the edge of the image were excluded. Lipid droplets were sorted into one of three size bins according to their area size: 0-50 μm^2^, 50-500 μm^2^, and >500 μm^2^, as used before for the purpose of lipid droplet size quantification.^14^ The lipid droplet occupied area within each size bin as a fraction of the total lipid droplet area was calculated for each image. The two acquired image per scaffold were averaged to obtain one data point per size bin for each scaffold.

### Gene expression analysis by RT-qPCR

RNA was isolated using the RNeasy Mini Kit (Qiagen). Scaffolds were washed with PBS and transferred to individual 1.5 mL microcentrifuge tubes with 350 μL RLT Lysis Buffer and 3.5 μL β-mercaptoethanol. Tubes were vigorously shaken using a vortex for 10 minutes at room temperature. RNA was extracted according to the manufacturer’s instructions and collected in 30 μL of distilled water. The RNA concentration was measured with a NanoDrop spectrophotometer (ThermoScientific). RNA was converted to cDNA by reverse transcription PCR with qScript cDNA Supermix (QuantaBio). 6 ng of cDNA was used per qPCR reaction consisting of gene-specific primers (Supplementary Information Table 3) and amplified with the PowerSYBR Green Master Mix (Applied Biosystems in a CFX-96 thermocycler (Bio-Rad). Data was acquired using the Bio-Rad CFX Manager 3.1. Gene expression was normalized to the geometric mean of the two reference genes *RPL13A* and *TBP*. Data was presented as the ΔΔCt value, relative to Day 0 of differentiation **(Figure 1d)** or to the vehicle control **(Figure 5c, Supplementary Information Figure 5, 7, 8, 10)**. The primers used are listed in **Supplementary Information Table 3**.

### Insulin sensitivity analysis

Scaffolds were serum-starved in DMEM/F12 for 4 hours and washed twice with PBS. To measure cell viability, the scaffolds were incubated in 50 μL CellTiter Fluor™ reagent (Promega) and 50 μL glucose-free DMEM/F12 (Gibco) for 1 hour at 37 °C. Fluorescence was measured using a plate reader (Infinite M Plex, Tecan) and excitation and emission wavelengths were set to 380/505 nm. Scaffolds were washed with PBS and incubated with or without 100 nM human insulin (Sigma-Aldrich) in 100 μL glucose-free DMEM/F12 for 1 hour at 37 °C. Scaffolds were washed with PBS and incubated with 50 μL 1 mM 2-deoxyglucose in PBS for 10 minutes at room temperature. Uptake of 2-deoxyglucose by the adipocytes was measured using a Glucose Uptake-Glo™ Assay kit (Promega) and luminescence was measured according to the manufacturer’s instructions using a plate reader. Luminescence values of uptake of 2-deoxyglucose were normalized by fluorescence values of cell viability.

### Lipolysis analysis

To measure cell viability, the scaffolds were incubated in 50 μL CellTiter Fluor™ reagent (Promega) and 50 μL glucose-free DMEM/F12 (Gibco) for 1 hour at 37 °C. Fluorescence was measured using a plate reader and excitation and emission wavelengths were set to 380/505 nm. To measure basal lipolysis, scaffolds were incubated in 100 μL 2 % fatty acid-free BSA (Sigma-Aldrich) in DMEM/F12 for 1 hour at 37 °C. Media was collected and basal rate of glycerol release by the adipocytes was measured using a Glycerol Glo™ Assay kit (Promega) and luminescence was measured according to the manufacturer’s instructions using a plate reader. To measure β-adrenergic-stimulated lipolysis, scaffolds were incubated in 100 μL 2 % fatty acid-free BSA with 10 μM isoproterenol-hydrochloride (Sigma-Aldrich) in DMEM/F12 for 1 hour at 37 °C. Media was collected and β-adrenergic-stimulated rate of glycerol release by the adipocytes was measured using a Glycerol Glo™ Assay kit and luminescence was measured according to the manufacturer’s instructions using a plate reader. Luminescence values of released glycerol were normalized by fluorescence values of cell viability.

### Cytokine analysis

On day 6 of fatty acid stimulation, the scaffolds were washed with PBS and incubated in 200 μL fresh DMEM/F12 supplemented with 1 % FBS. After 12 hours, conditioned media was collected in snap-cap tubes and stored at −80 °C until further processing. Once all samples were collected, the conditioned media of two of the same conditions within an experiment was combined to reach a volume of 400 μL. From this, 365 μL conditioned media was concentrated using an Amicon Ultra 10K Centrifuge Filter Unit as per the manufacturer’s instructions. Briefly, the samples were centrifuged at 14,000 *g* for 10 minutes at 4 °C to concentrate the proteins. The sample was recovered by inversely placing the filters in a snap-cap tube and centrifuging at 1000 *g* for 2 minutes at 4 °C, which yielded a volume of 75 μL (4.87x concentrated). Supernatant was filtered using a 1.2 μm and a 0.22 μm filter (Durapore) and cytokine levels were analyzed using the bead-based Human Cytokine Proinflammatory Focused 15-Plex Discovery Assay® Array (Eve Technologies, Alberta, Canada).

### Macrophage response to adipocyte conditioned media

Conditioned media was prepared on day 6 of fatty acid stimulation by washing the adipocyte scaffolds with PBS and incubating them in 300 μL of fresh RPMI-1640 supplemented with 10 % FBS and 1 % Penicillin-Streptomycin (Wisent Bioproducts). After 12 hours, conditioned media was collected and passed through a 0.22 μm 4 mm filter to remove particulates and debris while avoiding large volume loss. The conditioned media was stored at −80 °C until macrophage stimulation.

The collection and experimental use for the experiments demonstrated of primary human monocytes in **Figure 5c** and **Supplementary Information 10** was reviewed and approved by Canadian Blood Services Research Ethics Board (REB#2018.020) and The University of Toronto Office of Research Ethics reviewed the approved studies and further assigned administrative approval (REB#35956). Informed consent was obtained from all study participants. All methods in this study were performed in accordance with the guidelines and regulations of these two Research Ethics Boards. Human peripheral blood mononuclear cells (PBMCs) were isolated from the blood of healthy donors and differentiated to macrophages based on an established protocol.^29^ Briefly, PBMCs were isolated from blood using Ficoll-Paque (GE Healthcare) separation. The PBMC layer was collected, treated with red blood cell lysis buffer on ice for 10 minutes and PBMCs were collected by centrifugation at 400 *g* for 5 minutes and stored in liquid nitrogen. At the day of macrophage differentiation, PBMCs were thawed and passed through a 40 μm strainer to remove clumps of nonviable cells. PBMCs were centrifuged at 200 *g* for 5 minutes at 4 °C, collected and resuspended in 80μL of magnetic-activated cell sorting buffer (2 % FBS and 2 mM EDTA in calcium/magnesium-free PBS) per 10^7^ cells. CD14 microbeads (Miltenyi Biotec) were added to the cell suspension at 20 μL per 10^7^ cells and incubated for 15 minutes at 4 °C. Magnetic separation was performed using LS columns (Miltenyi Biotec) in a MACS separator in accordance with manufacturer’s instructions. For macrophage differentiation, CD14+ monocytes were plated in wells of a 24-well tissue culture plate at a density of 150,000 monocytes per well and cultured in macrophage differentiation media consisting of RPMI-1640 (Sigma-Aldrich) supplemented with 10 % FBS, 1 % Penicillin-Streptomycin, and M-CSF (Peprotech) at 50 ng/mL for 6 days. Macrophages were stimulated with conditioned media as described before.^20^ Briefly, after 6 days of macrophage differentiation, media was replaced with 270 μL of conditioned media from one adipocyte scaffold per well of macrophages and 90μL of fresh macrophage maintenance media consisting of RPMI-1640 supplemented with 10 % FBS and 1 % Penicillin-Streptomycin. After 48 hours, macrophages were lysed, and RNA extraction and qPCR were performed as described above.

### Statistical analysis

For most experiments, a minimum of three experimental replicates (N) were conducted each containing multiple technical replicate scaffolds (n); exception is the gene expression analysis in **Figure 1c** where two experimental replicates were performed. All graphs report the mean ± standard deviation of the mean (SD). If applicable, within a figure, data from technical replicate scaffolds within one experimental replicate are indicated by colour, where a colour indicates that the scaffolds were part of the same experimental replicate. GraphPad Prism software was used to conduct all statistical analysis presented. The statistical test, assumptions, and justifications, as well as the N and n for each statistical analysis are described in **Supplementary Information Table 4**. Significance is reported on all figures in star format with the associated p-values displayed in the corresponding figure legend.

## Results

### Optimization of human ADSC infiltration and differentiation in a 3D paper scaffold

We sought to establish a paper-based culture model of human adipose tissue to study phenotypic, morphological and functional changes associated with obesity. Towards this, our first goal was to devise a method to infiltrate and differentiate human adipose-derived stromal cells (hADSCs) into paper scaffolds to generate 3D mature adipocyte cultures. To do this we followed the experimental timeline as shown in **Figure 1a**. Briefly, hADSCs were suspended in a fibrin/Geltrex™ hydrogel and pipetted onto the surface of a dry paper scaffold (**Supplementary Information Figure 1**). The fibrin/Geltrex™ hydrogel formulation was selected based on our previous use for other non-diseased human cell types^21,30^. Upon contact with the paper surface, the hydrogel-cell mixture infiltrated in between the paper fibers of the highly porous scaffold via capillary wicking. Infiltrated scaffolds were then incubated at 37 °C to expedite fibrin polymerization. This produced a fiber-reinforced hydrogel with hADSCs evenly distributed throughout the gel in between the paper scaffold fibers **(Figure 1b)**.

After hydrogel gelation, scaffolds were incubated in growth media for two days after which the growth media was replaced by adipocyte differentation media for 14 days to induce specification of the hADSCs into mature adipocytes. To assess the effectiveness of this protocol to induce adipocyte differentiation in the hydrogel-infiltrated scaffolds, we assessed a variety of key adipocyte characteristics indicative of different maturity states. We first assessed morphology changes of the hydrogel embedded hADSCs as they differentiated into adipocytes within the scaffolds. As expected, during adipocyte differentiation, the elongated stromal cells underwent a morphological change towards a larger, spherical shape to accommodate the storage of lipids in spherical lipid droplets as observed using scanning electron microscopy **(Figure 1b)**. Further, we observed the formation of an ECM at day 14 suggesting that the differentiating adipocytes actively secrete ECM molecules within the three dimensional (3D) porous structure between scaffold fibers, demonstrating their interaction with the extracellular environment **(Figure 1b)**.

Another important feature associated with adipocyte differentiation is the formation of lipid droplets, which store energy in the form of triglycerides. To assess if lipid droplet formation occurred, we used BODIPY, a lipid-soluble dye, to stain cells in the scaffolds **(Figure 1c, Supplementary Information Figure 2)**. We observed large lipid droplets in adipocytes within the scaffolds on day 14 of differentiation. This is indicative of functional fatty acid uptake and triglyceride storage and suggested that human adipose-derived stromal cells cultured in the 3D paper scaffold maintain their expected capacity to differentiate into mature adipocytes as previously reported in 2D cultures.^11^ This phenotype was consistent across multiple donors.

We next set out to assess if the transcriptional differentiation dynamics of hADSCs differentiated into adipocytes in the 3D cultures were similar to those observed previously in 2D culture. We assessed the expression of adipocyte marker genes characteristic of key stages of adipocyte differentiation. Specifically, we quantified the expression of transcription factors peroxisome proliferator-activated receptor gamma (*PPARG*) and CCAAT/enhancer binding protein alpha (*CEBPA*), which upregulate expression of key adipocyte genes during differentiation.^31^ We also assessed the expression of downstream genes fatty acid translocase (*CD36*) and fatty acid binding protein 4 (*FABP4*) that facilitate lipid mobilization and storage, as well as downstream genes adiponectin (*ADIPOQ*) and leptin (*LEP*), two genes that encode for the characteristic hormones secreted by mature adipocytes.^9,32,33^ Our data suggested that changes in expression of each of these genes followed similar dynamics during adipocyte differentiation in the 3D paper scaffold as in standard 2D culture (**Figure 1d**). Taken together, these results suggest that our 3D hydrogel-infiltrated paper scaffold provides an environment compatible with human adipocyte differentiation and culture.

### Stimulation of the 3D adipocyte scaffolds with fatty acids induces an increase in lipid droplet size

Our next goal was to devise a treatment protocol to induce hallmarks of obese-like human adipocytes. We hypothesized that mimicking caloric overload in the 3D adipocyte scaffolds would induce aspects of an *in vivo* obese phenotype. To test this, we stimulated adipocyte-infiltrated scaffolds with a fatty acid-rich environment by adding either oleic acid, a C18:1 unsaturated fatty acid, or palmitic acid, a C16:0 saturated fatty acid, to the cell culture media **(Figure 2a)**. We selected these long-chain fatty acids because of their abundance in human plasma^34^ and their previous use in the literature.^14^ We explored the use of both fatty acids separately at multiple concentrations for various stimulation periods.

Obese adipocytes *in vivo* are characterized by an increased need for triglyceride storage in their lipid droplets, which corresponds to an increased lipid droplet size in *in vitro* cultures.^14^ Therefore, to assess the impact of each treatment we first analyzed lipid droplet size distributions using BODIPY staining in adipocytes in scaffolds treated for a period of 6 days with the different fatty acid-stimulation protocols **(Figure 2b).** We observed an increase in the abundance of large lipid droplets in the 3D adipocyte scaffolds treated with the fatty acid stimulation protocols compared to treatment with vehicle alone. To quantify this effect, we developed a FIJI workflow (**Supplementary Information Figure 4**) that classified all lipid droplets in the image into one of three size categories (0-50 μm^2^, 50-500 μm^2^, and >500 μm^2^). We then calculated the percentage of total lipid droplet-occupied area attributed to each of the three categories of lipid droplet size contained in each image. While we observed no changes in the contribution of lipid droplets with a size of 0-50 μm^2^ **(Figure 2c)**, we found a significant increase in the contribution of the largest lipid droplets (>500 μm^2^) in response to fatty acid stimulation. Of note, oleic acid appeared to show a dose-dependent increase in the fraction of large lipid droplets, however, palmitic acid did not produce dose-dependent effects. We also assessed the effect that the duration of fatty acid stimulation elicited on lipid droplet size changes in the 3D adipocyte scaffolds **(Supplementary Information Figure 3)** and concluded that lipid droplet hypertrophy appears to plateau at day 6 of fatty acid stimulation. Lipid droplets in adipocytes derived from a second primary cell line (line Z24.5) **(Supplementary Information Figure 2,3a)** consistently showed smaller lipid droplets in comparison to the P21.5 cell line **(Figure 1c, 2c)**. However, the trends in lipid droplet size increase were constant across multiple cell lines. These data suggest that stimulation of the 3D adipocyte scaffolds with both oleic acid and palmitic acid induces lipid droplet hypertrophy.

### Stimulation of the 3D adipocyte scaffolds with fatty acid impairs insulin sensitivity

Obese adipocytes *in vivo* that exhibit insulin resistance are thought to contribute to the development of metabolic disorder.^8^ To explore insulin resistance in fatty acid-treated adipocyte cultures, we first assessed the gene expression of enzymes involved in insulin signaling **(Supplementary Information Figure 5)**. Specifically, we analyzed the transcription of insulin receptor substrate 1 (*IRS1*) and Akt2 (*AKT2*), important enzymes in the insulin signaling pathway. Upon activation of this cascade, vesicles filled with insulin-dependent glucose transporter 4 (encoded by the *SLC2A4* gene) translocate from the cytosol and fuse with the plasma membrane to enable the uptake of glucose.^35^ We observed a trend towards decreasing gene expression of *IRS1* and *SLC2A4* in the oleic acid-treated adipocyte scaffolds, but not in those treated with palmitic acid, suggesting a fatty acid-specific effect on insulin signaling pathway gene expression.

To further assess the insulin sensitivity of our model, we employed a luminescence assay to analyze the uptake of glucose by the fatty acid-stimulated adipocytes in the scaffolds after incubation with insulin. The cellular mechanism of the assay is briefly illustrated in **Figure 3a**. In response to insulin, insulin-sensitive cells uptake more glucose than insulin-insensitive cells. As expected, we observed an increase in glucose uptake in the vehicle-stimulated 3D adipocyte scaffolds after incubation with insulin **(Figure 3b-d)**. However, we found that this increase in glucose uptake in response to insulin incubation was blunted in the 3D adipocyte scaffolds treated with either oleic acid or palmitic acid, independent of their concentration. These results suggest that treatment of the 3D adipocyte scaffolds with oleic acid or palmitic acid impairs insulin sensitivity.

### Stimulation of the 3D adipocyte scaffolds with palmitic acid, but not oleic acid, induces an increase in basal lipolysis

Previous work using animal models has demonstrated an increase in basal lipolysis and a decrease in β-adrenergic-stimulated lipolysis in obese adipocytes *in vivo*.^36^ While human obese adipocytes *ex vivo* also demonstrate an increased rate of basal lipolysis per cell number, they showed no attenuated β-adrenergic-stimulated lipolysis compared to lean cells.^37,38^ To determine if this increase in basal lipolysis after stimulation with fatty acids is observed at the level of gene expression, we analyzed the transcription of enzymes involved in lipolysis **(Supplementary Information Figure 7)**. We analyzed expression of adipose triglyceride lipase (*ATGL*), hormone sensitive lipase (*HSL*) and monoacylglyceride lipase (*MAGL*), genes that encode for enzymes involved in the sequential release of fatty acids from the glycerol backbone.^39^ With the exception of *ATGL* expression in the scaffolds stimulated with 500 μM palmitic acid, we found no clear trends in upregulation of these genes in the fatty acid-stimulated 3D adipocyte scaffolds compared to vehicle control.

We also assessed the lipolytic function of our fatty acid-stimulated 3D adipocyte scaffolds, by employing a functional assay that determines the release of glycerol from the adipocyte scaffolds as a measure for lipolysis, before and after stimulation with isoproterenol, a β-adrenergic stimulus **(Figure 4a)**. We observed that the 3D adipocyte scaffolds stimulated with the highest concentrations of palmitic acid demonstrated an increase in basal lipolysis compared to vehicle-stimulated scaffolds **(Figure 4b)**. While this increase in basal lipolysis was subtle, we observed a similar trend in increased basal glycerol release after oleic or palmitic acid stimulation in 3D scaffolds with adipocytes from a different donor **(Supplementary Information Figure 6a)**. However, after isoproterenol stimulation, we did not see a difference in lipolytic rate between the fatty acid-stimulated and the vehicle-stimulated 3D adipocyte scaffolds **(Figure 4c, Supplementary Information Figure 6b)**. Together, these results suggest that palmitic acid stimulation increases basal lipolysis, but does not alter β-adrenergic-stimulated lipolysis in the 3D adipocyte scaffolds.

### Conditioned media of palmitic acid-stimulated 3D adipocyte scaffolds alter gene expression of proinflammatory cytokines in macrophages

The obese adipose tissue microenvironment is characterized by inflammatory mediators such as lipids and cytokines that can affect macrophage phenotype *in vivo*.^40^ To investigate if our fatty acid-stimulated 3D adipocyte scaffolds could alter the inflammatory gene expression of macrophages, we performed an experiment using conditioned media from fatty acid-stimulated 3D adipocyte scaffolds as illustrated in **Figure 5a**. We observed that conditioned media from palmitic acid-stimulated 3D adipocyte scaffolds increased the expression of *IL6* and decreased the expression of *TNFA* in primary human macrophages. In contrast, the conditioned media from oleic acid-stimulated 3D adipocyte scaffolds had little effect on macrophage gene expression **(Figure 5b)**. Gene expression of inflammatory markers, *SPP1, IL1B* and *CCL2*, in macrophages was not affected by conditioned media from 3D adipocyte scaffolds stimulated with fatty acids compared to vehicle **(Supplementary Information Figure 9)**.

To investigate if the fatty acid-stimulated 3D adipocyte scaffold caused increased secretion of pro-inflammatory cytokines that could give rise to the change in macrophage phenotype, we performed a multiplex cytokine assay on the conditioned media. We observed high levels of cytokines that are known to be secreted by obese adipocytes, such as MCP1 and IL6. **(Figure 5c)** However, we did not find that fatty acid-stimulation of the 3D adipocyte scaffolds increased the expression of any of the cytokines that we analyzed. **(Figure 5c, Supplementary Information Figure 10)**. This suggests that the change in macrophage gene expression after incubation with conditioned media from fatty acid-stimulated 3D adipocyte scaffolds does not result from an increase in expression of common adipocyte-derived pro-inflammatory cytokines.

## Discussion

As a result of an increased need for lipid storage in obesity, the fat-storing adipocytes are one of the first cell types affected by caloric overload. Despite the advances in developing strategies to analyze whole adipose tissue dysfunction *in vitro*, no 3D model has yet focused on studying the unique contribution of dysfunctional adipocytes to metabolic disorder in obesity. We have developed a 3D *in vitro* human disease model that recapitulates functional characteristics of adipocytes during obesity, such as lipid droplet hypertrophy, insulin resistance, increased basal lipolysis and proinflammatory cytokine production.^41^

In our 3D model, we have exploited a biomaterial strategy that supports efficient adipocyte differentiation, demonstrated by the emergence of lipid droplets, which indicates functional lipid metabolism and triglyceride storage. Gene expression analysis demonstrated the upregulation of adipocyte transcription factors, lipid metabolism enzymes and essential adipokines throughout adipocyte differentiation. Notably, in line with previous studies, we found that during long-term culture of fatty acid stimulated adipocytes in a 2D monolayer, the cells detach from the tissue culture plastic as a result of large adipocyte size, high lipid content and spherical morphology.^11^ Our model offers an alternative to this by embedding the cells in a hydrogel and infiltrating this into a biocompatible scaffold. This was particularly valuable for allowing long-term 3D adipocyte culture in our application, since fatty acid stimulation further increases the lipid content and buoyancy of adipocytes. Importantly, the cellulose scaffold maintains its integrity throughout long-term culture, thereby providing a porous environment for the hydrogel-cell mixture to anchor to. Our work is also in line with previous studies demonstrating that several different biomaterial scaffolds and extracellular matrices can be used for effective adipocyte differentiation *in vitro*.^42–45^ Recently, trends in the development of self-assembled tissue cultures have sparked the creation of adipocyte spheroids cultures for the study of adipogenesis or adipocyte (dys)function.^16,46–48^ While these systems resemble physiologically relevant aspects of adipocyte biology, it is challenging to create specific tissue architectures using self-assembled aggregate cultures. Our strategy of infiltrating the hydrogel-cell mixture into a thin cellulose scaffold creates highly reproducible stackable ‘building blocks’ which opens the door for the investigation of endless combinations of possible tissue organizations. Further, the thin geometry of our model also enabled image-based analysis of obese adipocytes using conventional confocal microscopy.

Our assay mimics caloric overload by stimulating the 3D adipocyte scaffolds with oleic acid or palmitic acid, fatty acids that are abundantly present in human plasma,^34^ to induce important characteristics of dysfunctional adipocytes in obesity *in vivo*. Specifically, we found that both fatty acids were able to induce lipid droplet hypertrophy and insulin resistance in the 3D adipocyte scaffolds, while only palmitic acid was able to increase adipocyte basal lipolysis and induce an adipocyte secretome capable of altering macrophage phenotype. Importantly, our results indicate that gene expression analysis does not correlate with the functional effects of fatty acid stimulation on adipocytes with regards to insulin resistance, basal lipolysis and cytokine release profile. This demonstrates that employing functional assays to assess the downstream effects of caloric overload in adipocytes is essential for the characterization of the adipocyte phenotype as opposed to analyzing the cells on a transcriptional level alone. We also found that fatty acid stimulation increased functional basal lipolysis, but did not impair lipolysis in response to isoproterenol. This is in line with previous studies that have observed that freshly isolated human adipocytes from obese subjects compared to non-obese subjects demonstrated an increased basal lipolysis and a similar lipolytic rate upon β-adrenergic stimulation when normalized to cell number.^37,38^ In addition, we demonstrated that stimulating macrophages with the conditioned media from palmitic acid-stimulated 3D adipocyte scaffolds altered the expression of *IL6* and *TNFA*. This suggests a paracrine effect of palmitic acid-stimulated adipocytes on macrophage phenotype. While previous studies using 2D *in vitro* models have found that palmitic acid induces a pro-inflammatory cytokine secretion profile in adipocytes,^14,49^ we did not observe an effect of either oleic or palmitic acid stimulation on secretion of important pro-inflammatory cytokines by the adipocyte scaffolds. This suggests that the effect of conditioned media of palmitic acid-stimulated 3D adipocyte scaffolds on macrophage phenotype might be caused by an increase in lipolysis rather than an increase in pro-inflammatory cytokine secretion. Indeed, previous work has found that fatty acid-stimulation of macrophages alone, induces an pro-inflammatory gene expression profile.^50,51^ Other studies describe adding a mixture of different lipids to a vascularized adipocyte spheroid model to induce lipid droplet hypertrophy, an altered cytokine secretion profile, insulin sensitivity and lipolysis.^16^ However, our work suggests that palmitic acid stimulation mimics features of caloric overload-induced adipocyte dysfunction, without the need for interaction with other adipose tissue cell types.

Various studies have demonstrated that the ECM of adipose tissue mainly consists of fibrillar collagens (mainly type I and IV), laminins, heparan sulphate proteoglycans, and fibronectin,^52–54^ with the latter drastically decreasing during adipocyte differentiation.^54,55^ Other groups have used various extracellular matrices for adipose tissue engineering, such as collagen I^56–58^ or an alternative basement membrane matrix.^59,60^ In order to further recapitulate features of the adipose tissue microenvironment, it may be necessary to adjust the composition of the hydrogel to reflect the dominance of collagen proteins in adipose tissue. This was not essential for any of the metrics studied here but may be an important parameter to study other aspects of metabolic disease in adipocytes. Given the modularity of our engineered adipocyte platform, it can easily be adjusted to explore extracellular matrices with diverse chemical and mechanical properties.

We chose to differentiate human adipose-derived stromal cells to adipocytes in our 3D scaffold as opposed to infiltrating mature human adipocytes in the scaffold. This strategy provided us with a continuing supply of proliferating stromal cells instead of relying on a finite number of terminally differentiated mature adipocytes that were freshly isolated from human material. The limitation of this approach however is that *in vitro* adipogenesis yields adipocytes that do not fully resemble the mature adipocyte phenotype *in vivo*. For example, adipocytes differentiated *in vitro* generally contain multiple smaller lipid droplets, while mature adipocytes *in vivo* are characterized by a single, large lipid droplet.^61^ Other groups have integrated mature human adipocytes in 3D models and have demonstrated that the adipocytes maintain their phenotype a unilocular lipid droplet.^62,63^ By using these strategies, future experiments can incorporate mature human adipocytes from patients with obesity or metabolic disorder to investigate the cellular mechanisms that are altered in these adipocytes. While we observed constant trends in the effect of fatty acid stimulation on lipid droplet hypertrophy and increased basal lipolysis, we did find variability in the magnitude of the effect between different primary cell lines. This suggests that donor-to-donor variability is reflected in the ADSC-derived adipocytes which can be employed in the future for the purpose of patient stratification.

While our model enables the investigation of the unique contribution of obese adipocytes to metabolic disorder, it currently does not incorporate any other adipose tissue cell types. Several characteristics of obese adipose tissue are attributed to the interaction between adipocytes and other cell types. For example, previous work has suggested that chronic inflammatory signalling from immune cells in the obese adipose tissue attenuates β-adrenergic signaling in adipocytes.^64^ As our model consists of adipocytes alone, this could explain why impaired β-adrenergic signaling was not recapitulated in our functional lipolysis data. Incorporating other adipose tissue cell types into our model may further enhance the recapitulation of the *in vivo* obese adipose tissue phenotype. Other groups have created vascularized adipocyte models incorporating human adipocytes with endothelial cells,^16,48,56,65^ immune cells^66^ or cancer cells^67,68^ to understand the interaction between adipocytes and these various cell types. The geometry and modularity of our scaffold platform is well suited for incorporating additional cell types in future studies. It will be important however to only introduce the minimal complexity required to capture the biology of interest to avoid over complicating the culture assay.

## Conclusion

In this study, we describe the development and functional validation of a human 3D *in vitro* disease model of obese adipocytes, using a thin cellulose scaffold-based system that allows for easy tissue architecture control and image-based analysis. To induce an obese phenotype, we mimicked caloric overload by stimulating the 3D adipocyte scaffolds with oleic or palmitic acid. We observed that the effect of palmitic acid treatment increased lipid droplet size, insulin resistance, basal lipolytic rate and macrophage activation. This model can be employed to study the unique mechanisms by which adipocyte dysfunction contributes to metabolic disorder in obesity.

## Supporting information

Pieters et al_Supplementary Information

## Acknowledgements

The authors would like to thank Audrey Darabie for her advice on SEM optimization and Ileana Co and Aleksandra Fomina for providing the macrophages for the adipocyte-conditioned media experiments.

## Author contributions

Conceptualization: PMG, APM, VMP

Formal analysis: VMP, STR, PMG, APM

Funding acquisition: PMG, APM

Investigation: VMP, STR, SK, NTL

Methodology: VMP, KS, STR, NTL, SK, SNV

Project administration: VMP, PMG, APM

Resources: ADC, KS

Software: STR, VMP

Supervision: PMG, APM, VMP

Validation (Software): STR

Visualization: VMP

Writing - Original Draft: PMG, APM, VMP, STR, KS, NTL

Writing - review & editing: All authors

## Data and materials availability

The datasets generated during and analyzed in this study are available from the corresponding author on reasonable request.

