## Supplementary material for "A three-dimensional human adipocyte model of fatty acid-induced obesity": Pieters et al_Supplementary Information

**Supplementary Information Figure 1. Scanning electron microscopy image of paper scaffold**. **a.** Representative scanning electron microscopy image of dry paper scaffold, not infiltrated with hydrogel or cells. Magnification 100x; scale bar, 100 µm. **b.** Representative scanning electron microscopy image of paper scaffold infiltrated with the fibrin/Geltrex™ hydrogel, without cells. Magnification 100x; scale bar 100 µm.

**a**

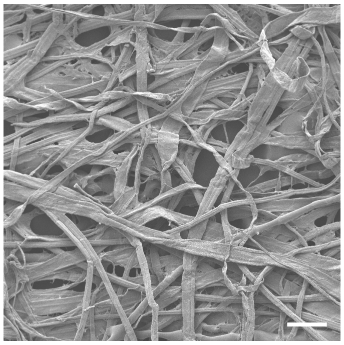

**b**

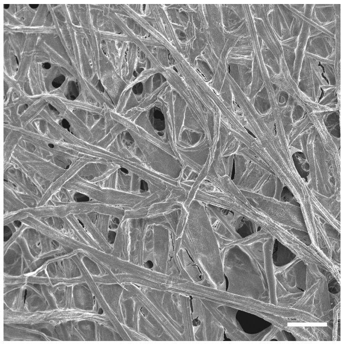

**Supplementary Information Figure 2. Adipocyte differentiation in 3D paper scaffold using human adipose-derived stromal cells derived from an additional donor.** Representative confocal images of 3D paper scaffold with human adipose-derived stromal cells at 0 and 14 days of differentiation derived from an additional donor. The orthogonal view of the respective scaffold is displayed below. Scaffolds were stained with the lipid-soluble dye BODIPY (green) and DNA stain DRAQ5 (magenta). Magnification 20x; scale bar, 50 μm.

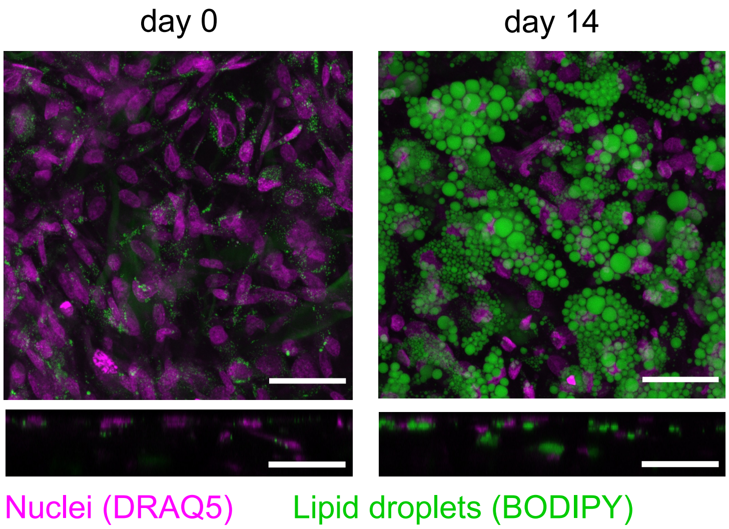

**Supplementary Information Figure 3. Fatty acid stimulation of human adipocytes in 3D paper scaffold increases lipid droplet size in a duration-dependant manner**. **a.** Quantification of lipid droplet size in adipocytes in 3D paper scaffold stimulated with oleic acid or palmitic acid in various concentrations for 4 days. Lipid droplets were grouped in different size categories of 0-50 µm2 (light grey), 50-500 µm2 (medium grey), and >500 µm2 (dark grey). Data is displayed as the percentage of lipid droplet area that is occupied by lipid droplets in the three different size categories. Experiments were performed with cells derived from one human donor. Each data point represents the average of two images of a 3D paper scaffold. Error bars represent standard deviation. n=2 to 3 scaffolds/stimulation condition. **b.** Quantification of lipid droplet size in adipocytes in 3D paper scaffold stimulated with oleic acid or palmitic acid in various concentrations for 8 days. Lipid droplets were grouped in different size categories of 0-50 µm2 (light grey), 50-500 µm2 (medium grey), and >500 µm2 (dark grey). Data is displayed as the percentage of lipid droplet area that is occupied by lipid droplets in the three different size categories. Experiments were performed with cells derived from one human donor. Each data point represents the average of two images of a 3D paper scaffold. Error bars represent standard deviation. n=3 scaffolds/stimulation condition. Statistics performed by one-way ANOVA and post-hoc Dunnet's test, comparing the lipid droplet-occupied area in the >500 µm2 size category of each fatty acid stimulated group to the vehicle control. *p <0.05; **p<0.01.

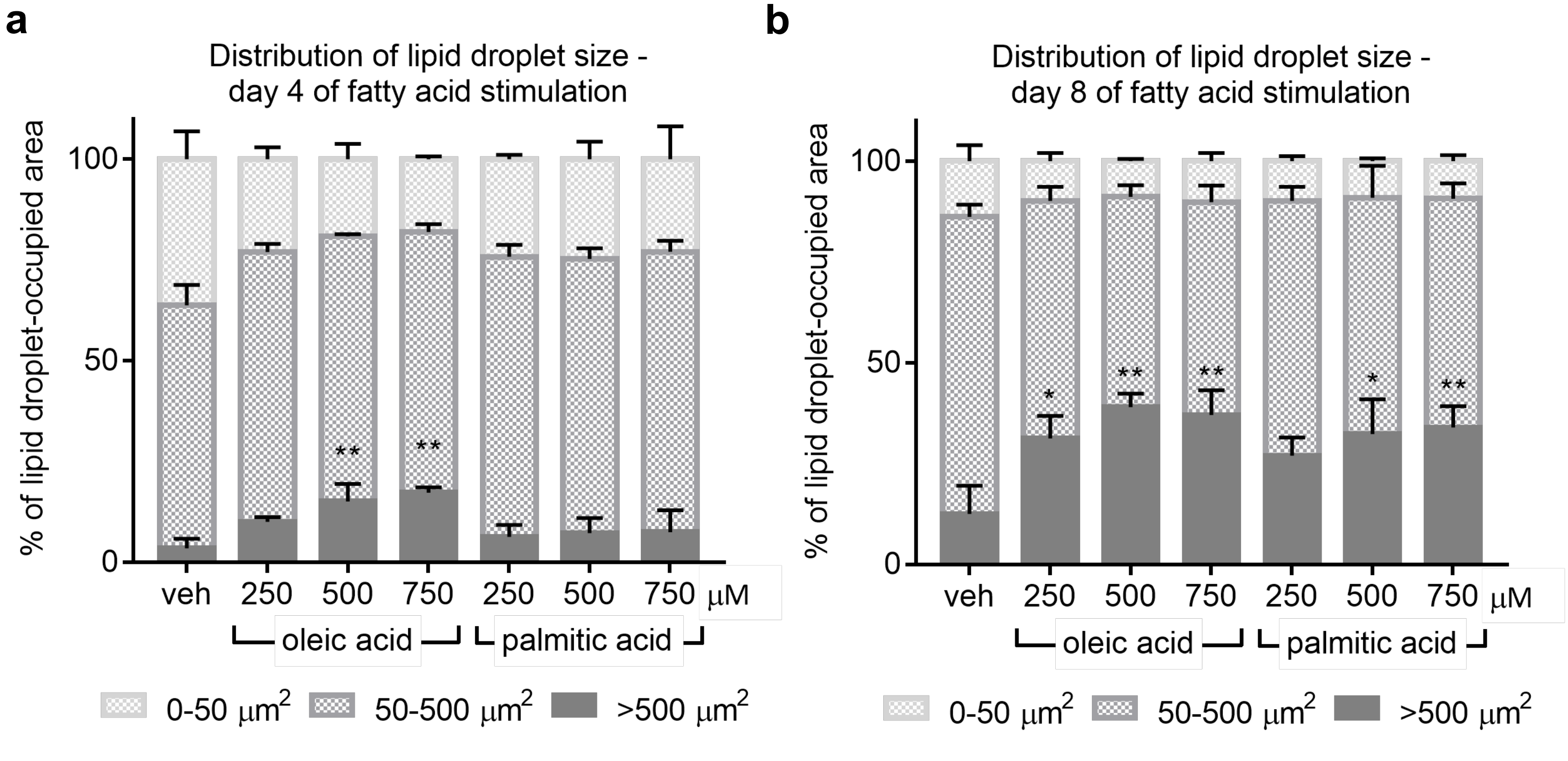

**Supplementary Information Figure 4. Image analysis workflow developed to quantify lipid droplet size in 3D adipocyte scaffold. a.** Illustration of the image analysis workflow steps. First, 3D confocal stacks are converted to a 2D image using a Maximum Intensity Projection. The projected image was sharpened using a 3x3 filter kernel with centre element "15" and all other elements "-1" and subsequently binarized using FIJI's Moments thresholding algorithm. The resulting objects were segmented using FIJI's Watershed tool, followed by three iterations of Openings (count=7) to clean up any background pixels, jagged edges, or holes in lipid droplets stemming from upstream processing. The finalised mask was fed to FIJI's Analyze Particles tool where the area of each object and total number of objects in the image were extracted. Objects on the edge were excluded. **b.** Validation of image analysis workflow by comparison to manual annotation. Each data point represents the lipid droplet-occupied area of >500 μm² lipid droplets of an image quadrant quantified by manual annotation and by the image analysis workflow. The black line represents the linear regression line of the data points and the R2 value indicates the coefficient of determination.

**a**

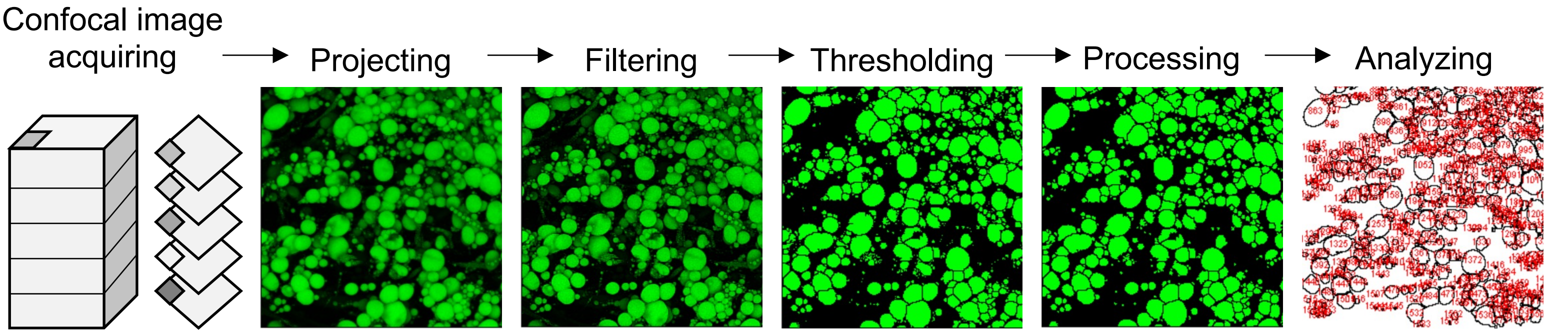


**b**

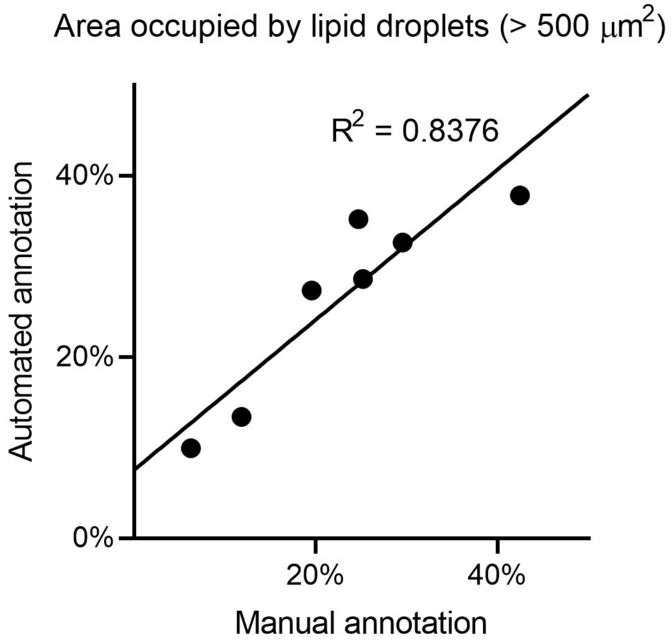

**Supplementary Information Figure 5. Fatty acid stimulation of human adipocytes in 3D paper scaffold alters gene expression of insulin-signaling related genes.** Three experiments were performed with adipocytes derived from one human donor, identifiable by the light grey, dark grey or black data points. Data is displayed as ΔΔCt relative to vehicle, using the geometric mean of reference genes *RPL13A* and *TBP*. Each data point represents the average of three RT-qPCR repeated measures. Error bars represent standard deviation. n=4 for 750 µM PA, 9 for 750 µM OA and 6 for all other groups. Statistics performed by one-way ANOVA and post-hoc Dunnet's test. *p <0.05; **p<0.01; ***p<0.001.

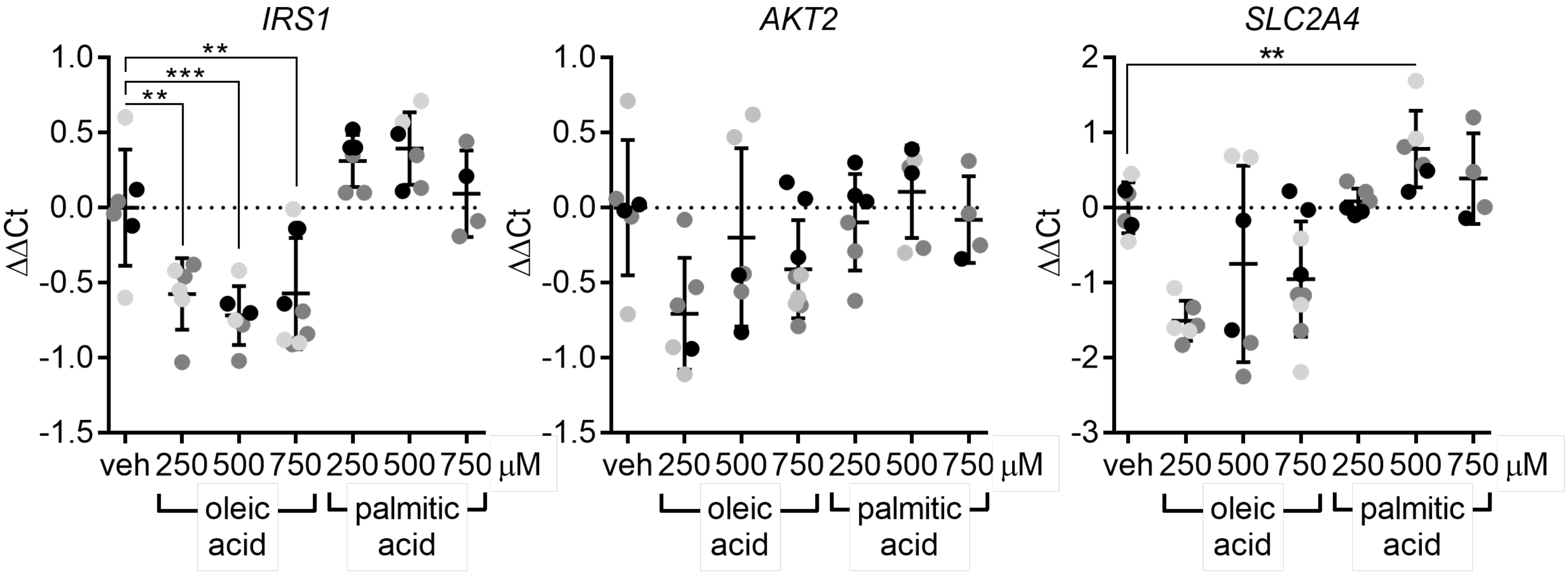

**Supplementary Information Figure 6. Fatty acid stimulation of human adipocytes from a different donor in 3D paper scaffold increases lipolysis. a.** Luminescence assessment of basal glycerol release of 3D adipocyte scaffolds after stimulation with various concentrations of oleic or palmitic acid. Data is displayed as concentration of glycerol released by the cells in each scaffold, normalized by a fluorescent measurement of cell viability of each scaffold. One experiment was performed with cells from one human donor. Each data point represents a 3D adipocyte scaffold. **b.** Luminescence assessment of isoproterenol-incubated glycerol release of 3D adipocyte scaffolds after stimulation with various concentrations of oleic or palmitic acid. Data is displayed as concentration of glycerol released by the cells in each scaffold, normalized by a fluorescent measurement of cell viability of each scaffold. One experiment was performed with cells from one human donor. Each data point represents a 3D adipocyte scaffold. Statistics performed by one-way ANOVA and post-hoc Dunnet's test. *p <0.05.

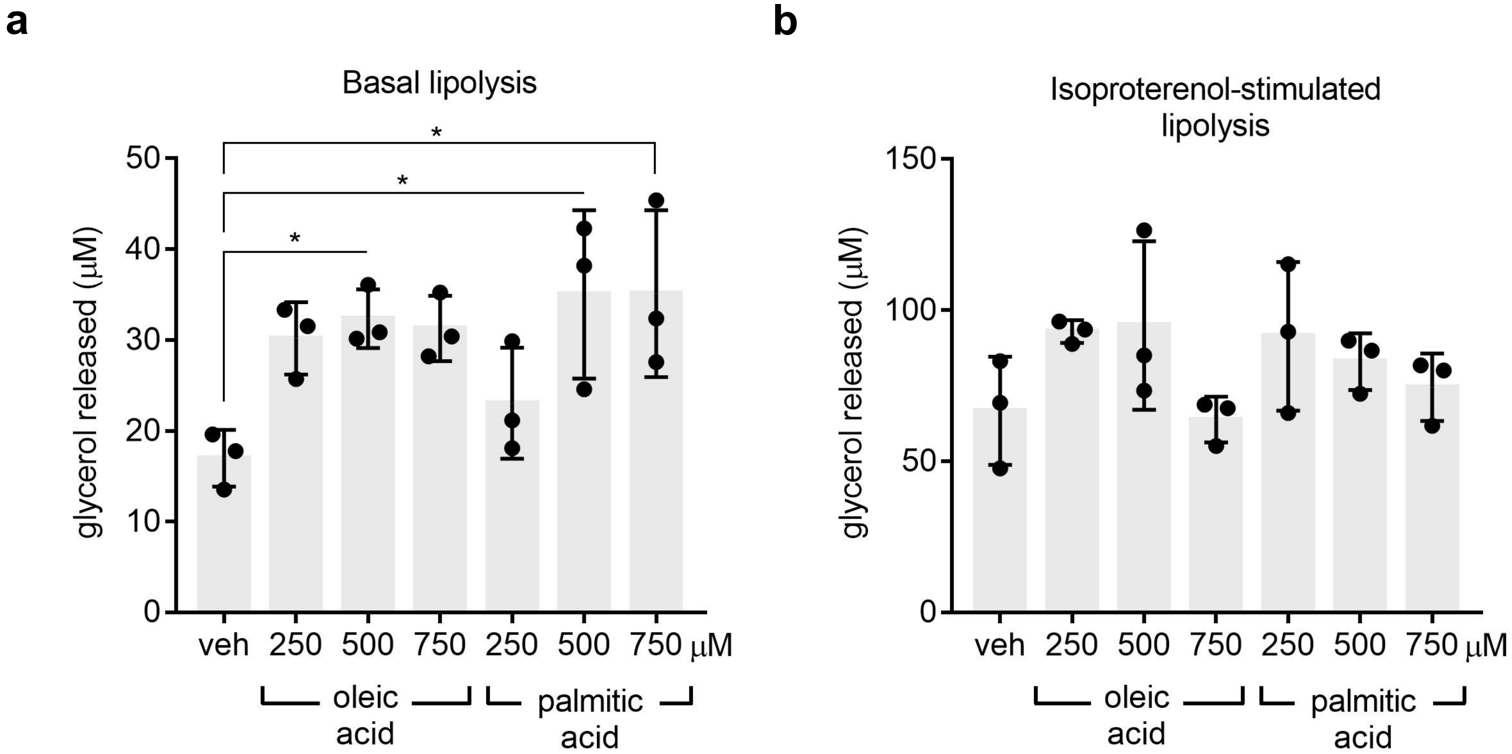

**Supplementary Information Figure 7. Fatty acid stimulation of human adipocytes in 3D paper scaffold alters gene expression of lipolysis-related genes.** Three experiments were performed with adipocytes derived from one human donor, identifiable by the light grey, dark grey or black data points. Data is displayed as ΔΔCt relative to vehicle, using the geometric mean of reference genes *RPL13A* and *TBP*. Each data point represents the average of three RT-qPCR repeated measures. Error bars represent standard deviation. n= 6 for vehicle, 500 μM OA and 500 μM PA, 7 for 750 μM PA and 9 for all other groups. Statistics performed by one-way ANOVA and post-hoc Dunnet's test. *p <0.05; **p<0.01.

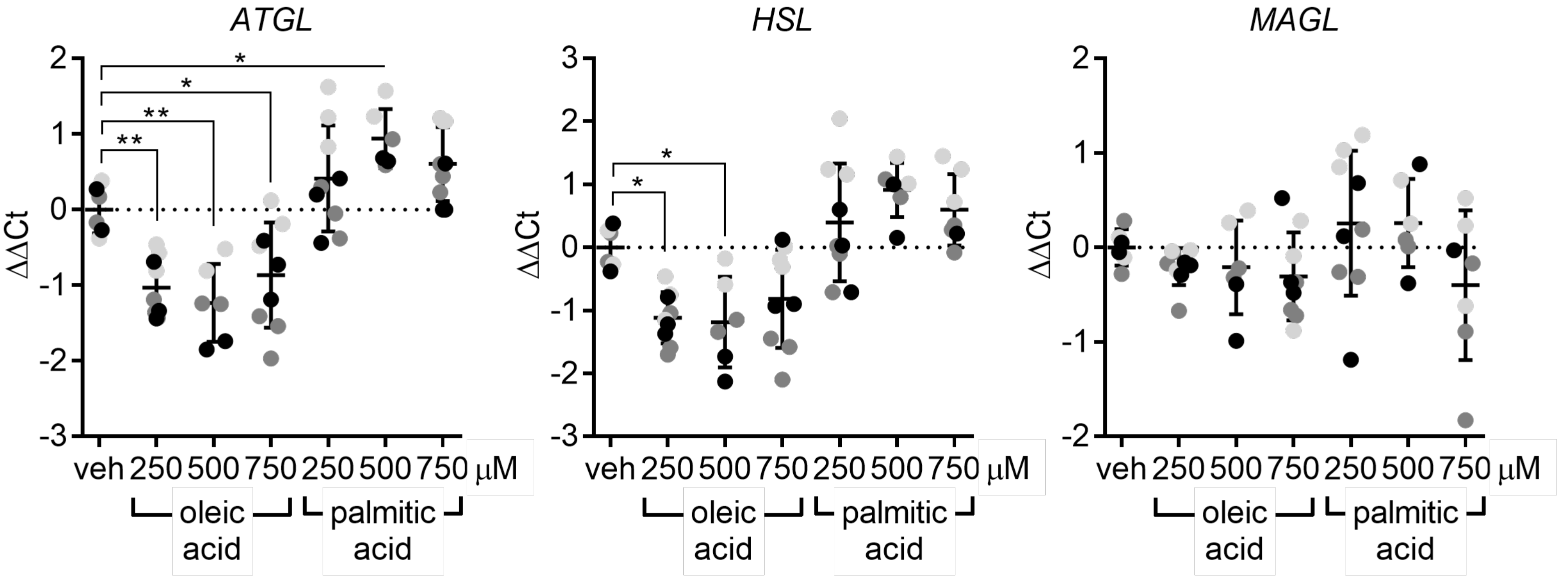

**Supplementary Information Figure 8. Fatty acid stimulation of human adipocytes in 3D paper scaffold alters expression of cytokine genes.** Three experiments were performed with adipocytes derived from one human donor, identifiable by the light grey, dark grey or black data points. Data is displayed as ΔΔCt relative to vehicle, using the geometric mean of reference genes *RPL13A* and *TBP*. Each data point represents the average of three RT-qPCR repeated measures. Error bars represent standard deviation. n= 6 for vehicle, 500 μM OA and 500 μM PA, 7 for 750 μM PA and 9 for all other groups. Statistics performed by one-way ANOVA and post-hoc Dunnet's test. *$p < 0.05$; **$p<0.01$; ***$p<0.001$; ****$p<0.0001$.

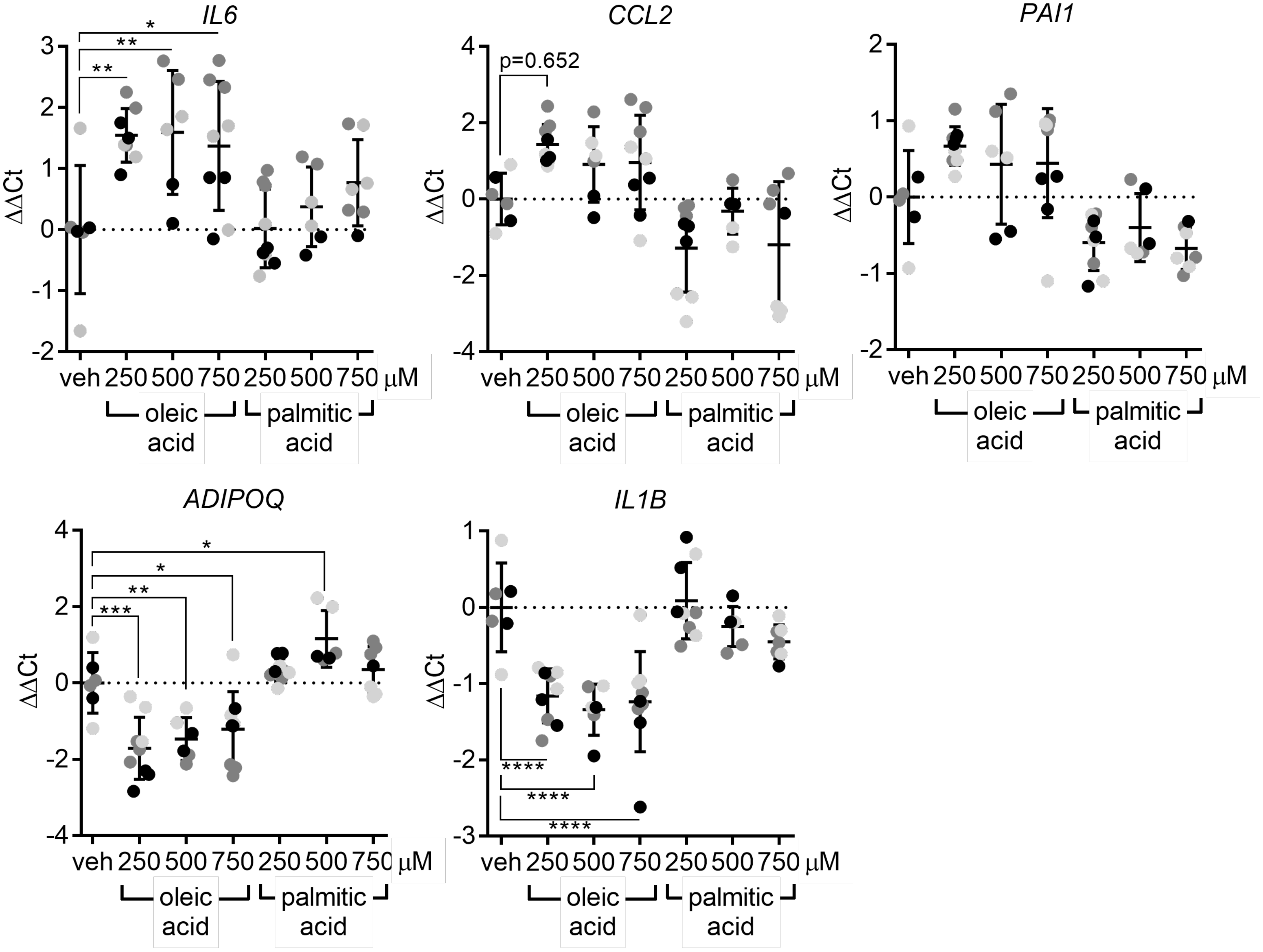

**Supplementary Information Figure 9. Conditioned media from fatty acid-stimulated 3D adipocyte scaffolds affects gene expression of certain pro-inflammatory markers in human macrophages**. RT-qPCR analysis of pro-inflammatory gene expression in macrophages in response to stimulation with adipocyte-conditioned media. Three experiments were performed with adipocyte-conditioned media from one human donor, identifiable by the green, orange or blue data points. Data is displayed as ΔΔCt relative to vehicle, using the geometric mean of reference genes *RPL13A* and *TBP*. Experiments were performed with cells derived from one human macrophage donor. Each data point represents the average of three RT-qPCR technical replicates of one 2D well of macrophages, stimulated with conditioned media from one 3D paper scaffold. Error bars represent standard deviation. n=9 (for vehicle) or 10 (for fatty-acid stimulated adipocyte conditioned media). Statistics performed by one-way ANOVA and post-hoc Dunnet's test.

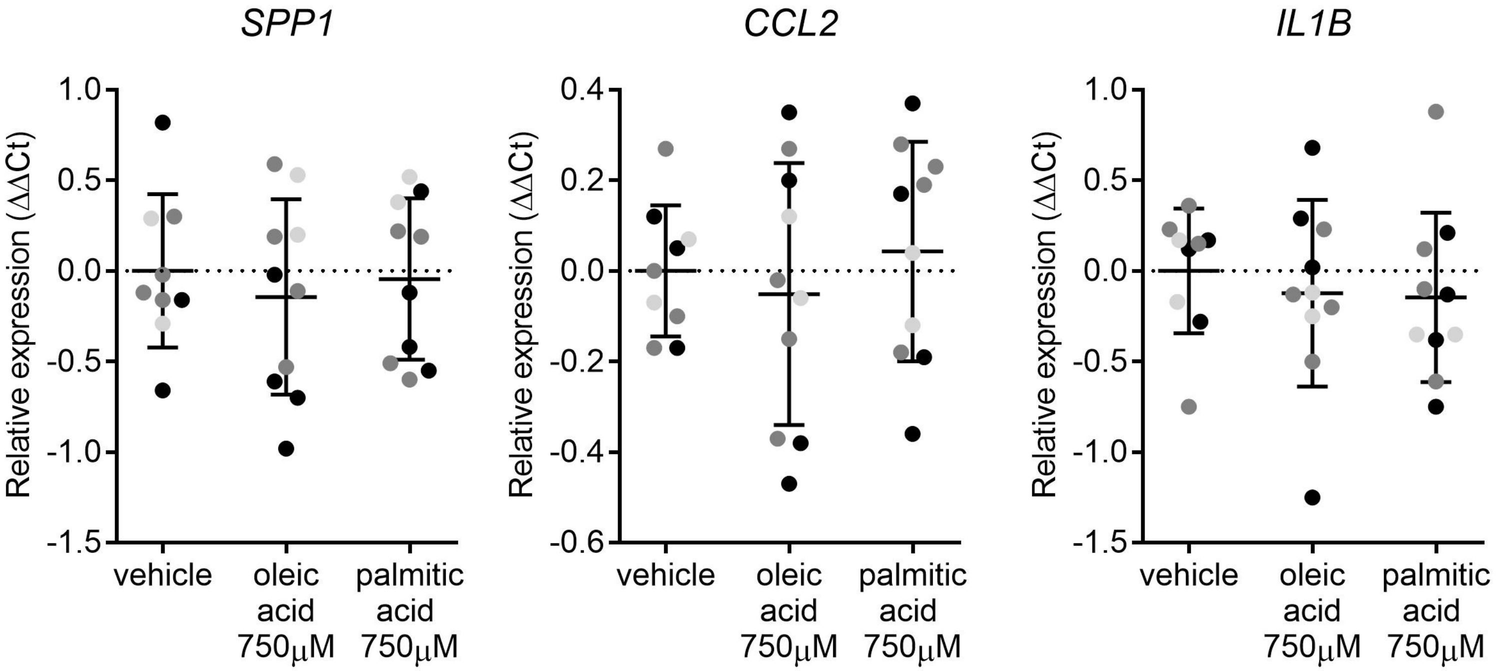

**Supplementary Information Figure 10. Fatty acid stimulation of human adipocytes in 3D paper scaffold does not affect secretion of pro-inflammatory cytokines**. Multiplex analysis of cytokines released into the cell culture media from 3D adipocyte scaffolds in response to stimulation with fatty acids. Three experiments were performed with adipocytes derived from one human donor, identifiable by the light grey, dark grey or black data points. Data is displayed as the concentration of released cytokines into the cell culture media (pg/mL). Each data point represents the average of two measurements of the cell culture media from one 3D adipocyte scaffold. Error bars represent standard deviation. n= 6. Statistics performed by one-way ANOVA and post-hoc Dunnet's test, no significance p values were observed.

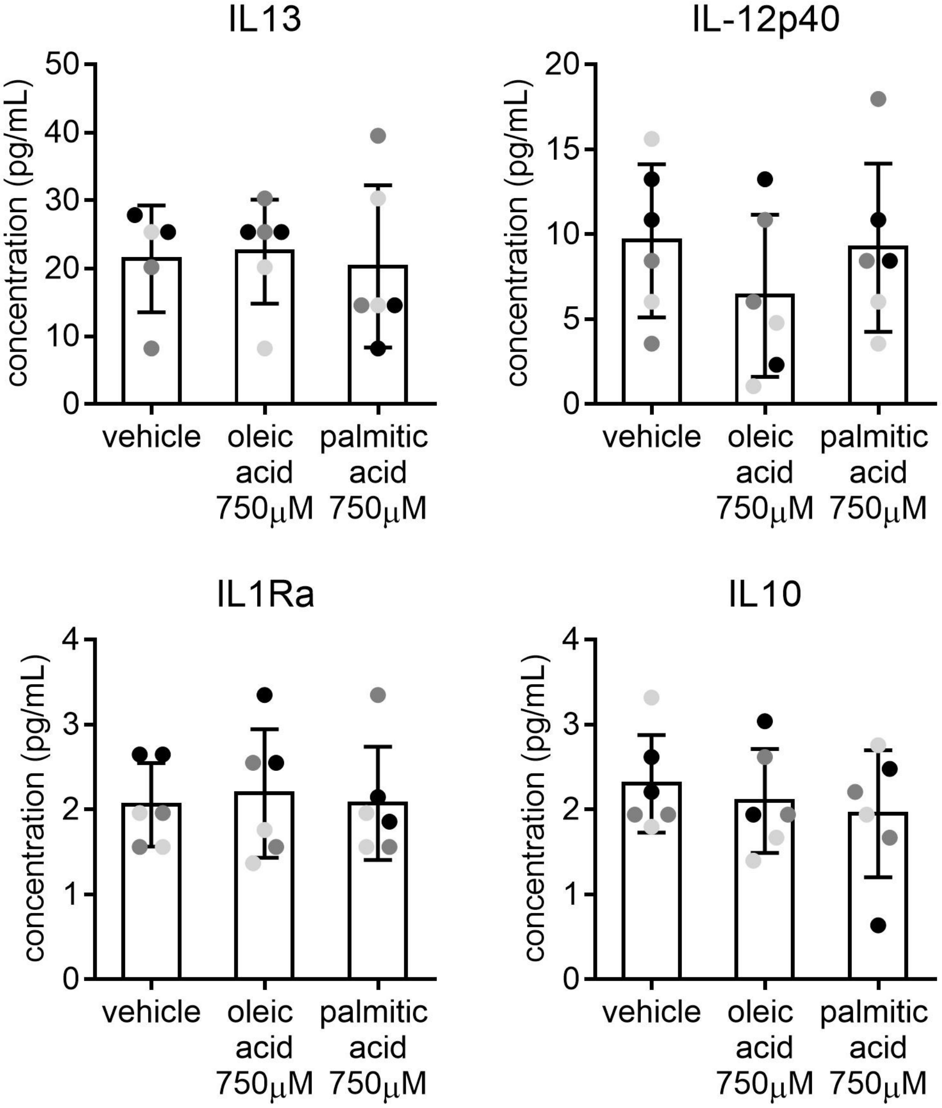

**Supplementary Information Table 1. Adipose-derived stromal cell donor information.**

| Cell ID number | Source | LOT# | Donor age (years) | Donor sex | Donor BMI | Adipose Tissue Depot | Used in figure |
|---|---|---|---|---|---|---|---|
| P21.5 | Dr. Arianna Dal Cin | N/A | 38 | Female | 21.5 | Breast | 1b, 1c, 2, 3, 4, 5, 6, SI1, SI3b, SI5, SI7, SI8, SI9 |
| Z25.3 | ZenBio | ASC01250 | 40 | Male | 25.3 | Abdomen | 1d |
| Z24.5 | ZenBio | ASC03070 | 49 | Male | 24.5 | Thigh/Hip | 1d, 2c, SI2, SI3a, SI6 |

### Supplementary Information Table 2. Composition of media and solutions used.

| Solution | Reagent | Manufacturer | CAT # |
|---|---|---|---|
| ADSC growth medium | DMEM/F12 | Gibco | 11320033 |
| | 10% FBS | Wisent Bioproducts | 098-150 |
| | 10ng/mL hbFGF | Peprotech | 100-18B |
| Adipocyte differentiation medium | DMEM/F12 | Gibco | 11320033 |
| | 10% FBS | Wisent Bioproducts | 098-150 |
| | 250μM IBMX | Sigma-Aldrich | I5879 |
| | 100μM indomethacin | Sigma-Aldrich | I7378 |
| | 0.5μM dexamethasone | Sigma-Aldrich | D2915 |
| | 10mg/ml human insulin | Sigma-Aldrich | I9278 |
| Fibrinogen solution | 0.9 % wt/vol NaCl in distilled water | N/A | N/A |
| | 10 mg/mL fibrinogen | Sigma-Aldrich | F8630 |
| ECM master mix | 40% DMEM/F12 | Gibco | 11320033 |
| | 40% 10mg/mL fibrinogen | N/A | N/A |
| | 20% Geltrex™ | Gibco | LSA1413301 |
| Oleic acid media | DMEM/F12 | Gibco | 11320033 |
| | 1% FBS | Wisent Bioproducts | 098-150 |
| | 250μM/500μM/750μM BSA-conjugated oleate | Cayman Chemicals | 29557-5 |
| Palmitic acid media | DMEM/F12 | Gibco | 11320033 |
| | 1% FBS | Wisent Bioproducts | 098-150 |
| | 250μM/500μM/750μM BSA-conjugated palmitate | Cayman Chemicals | 29558-5 |
| Vehicle media | DMEM/F12 | Gibco | 11320033 |
| | 1% FBS | Wisent Bioproducts | 098-150 |
| | 80μM fatty acid-free BSA | Sigma-Aldrich | A8806 |

**Supplementary Information Table 3. Sequences of qPCR primers used.**

| Gene | Forward primer | Reverse primer |
|---|---|---|
| *ADIPOQ* | GGTGAGAAGGGTGAGAAAGGA | TTTCACCGATGTCTCCCTTAG |
| *AKT2* | CTCACACAGTCACCGAGAGC | TGGGTCTGGAAGGCATACTT |
| *ATGL* | ACCAGCATCCAGTTCAACCT | ATCCCTGCTTGCACATCTCT |
| *CCL2* | AGTCTCTGCCGCCCTTCT | GTGACTGGGGCATTGATTG |
| *CD36* | CCTGGCTGTGTTTGGAGGTA | AATGAGAAGGCATATTCTTTGGAC |
| *CEPBA* | GGAGCAAATCGTGCCTTGTC | CTTCTCTCATGGGGGTCTGC |
| *FABP4* | GATAAACTGGTGGTGGAATGCG | ATGCGAACTTCAGTCCAGGT |
| *HSL* | GAGTTAAGTGGGCGCAAGTC | AAGTCCCTCAGGGTCAGGTT |
| *IL1B* | AATCTGTACCTGTCCTGCGTGTT | TGGGTAATTTTTGGGATCTACACTCT |
| *IL6* | CCAGTACCCCCAGGAGAAGATT | CATGTCTCCTTTCTCAGGGCTG |
| *IRS1* | TATGCCAGCATCAGTTTCCA | TTGCTGAGGTCATTTAGGTCTTC |
| *LEP* | TCCCCTCTTGACCCATCTC | GGGAACCTTGTTCTGGTCAT |
| *MAGL* | TCTGACTTCCACGTTTTCGTC | AGAACCAGAGGCGAAATGAGT |
| *PAI1* | CCTCAGGAAGCCCCTAGA | TGGAGAGGCTCTTGGTCT |
| *PPARG* | AAGGGGCCTTAACCTCTGCT | GGTCATTTCGTTAAAGGCTGACTC |
| *RPL13A* | CATAGGAAGCTGGGAGCAAG | GCCCTCCAATCAGTCTTCTG |
| *SLC2A4* | GAGCAGGACAGGAGACAAGAA | AGAGTCTGCGTGGCAAGAAT |
| *SPP1* | GCCGAGGTGATAGTGTGGTT | AACGGGGATCGGCCTTGTATG |
| *TBP* | GCCGGCTGTTTAACTTCGC | CAAGAAACAGTGATGCTGGGTC |
| *TNFA* | TCAGATCATCTTCTCGAACCCC | ATCTCTCAGCTCCACGCCAT |

**Supplementary Information Table 4. Statistical test, assumptions, and justifications.**

| Fig. | Biological Experiment (N) | Total scaffolds per condition (n) | Repeated measure | n for stats/ error bars | Statistical test | Test assumptions | Justification |
|---|---|---|---|---|---|---|---|
| **1d** | 2 | 6 | 3 | 6 | Two-sample t-test with Bonferroni correction | Approximate normality | Shapiro-Wilk (normality), Brown-Forsythe (eq. var.) |
| **2c** | 3 | veh, 250 OA, 750 OA, 750 PA (8); 500 OA, 250 PA (7); 500 PA (4) | 2 | 4, 7 or 8 | One-way ANOVA, post-hoc Dunnett's | Constant variance and approximate normality | Shapiro-Wilk (normality), Brown-Forsythe (eq. var.) |
| **3** | 3 | insulin, basal vehicle (8); basal OA, basal PA (7) | 1 | 7 or 8 | Two-way ANOVA, post-hoc Sidak's | Constant variance and approximate normality | Shapiro-Wilk (normality), Brown-Forsythe (eq. var.) |
| **4** | 3 | 10 | 1 | 10 | One-way ANOVA, post-hoc Dunnett's | Constant variance and approximate normality | Shapiro-Wilk (normality), Brown-Forsythe (eq. var.) |
| **5b** | 3 (1x macrophage stimulation) | 10 | 3 | 10 | One-way ANOVA, post-hoc Dunnett's | Constant variance and approximate normality | Shapiro-Wilk (normality), Brown-Forsythe (eq. var.) |
| **5c** | 3 | 6 | 2 | 6 | One-way ANOVA, post-hoc Dunnett's | Constant variance and approximate normality | Shapiro-Wilk (normality), Brown-Forsythe (eq. var.) |
| **SI 3** | 1 | 250 OA, 500 OA, 250 PA, 500 PA (3); veh, 750 OA, 750 PA (2) | 2 | 2 or 3 | One-way ANOVA, post-hoc Dunnett's | Constant variance and approximate normality | Shapiro-Wilk (normality), Brown-Forsythe (eq. var.) |
| **SI 5** | 3 | 750 OA (9); veh, 250 OA, 500 OA, 250 PA, 500 PA (6); 750 PA (4) | 3 | 4, 6 or 9 | One-way ANOVA, post-hoc Dunnett's | Constant variance and approximate normality | Shapiro-Wilk (normality), Brown-Forsythe (eq. var.) |
| **SI 6** | 1 | 3 | 1 | 3 | One-way ANOVA, post-hoc Dunnett's | Constant variance and approximate normality | Shapiro-Wilk (normality), Brown-Forsythe (eq. var.) |
| **SI 7** | 3 | 250 OA, 750 OA, 250 PA, (9); 750 PA (7); veh, 500 OA, 500 PA (6) | 3 | 6, 7 or 9 | One-way ANOVA, post-hoc Dunnett's | Constant variance and approximate normality | Shapiro-Wilk (normality), Brown-Forsythe (eq. var.) |
| **SI 8** | 3 | 250 OA, 750 OA, 250 PA, (9); 750 PA (7); veh, 500 OA, 500 PA (6) | 3 | 6, 7 or 9 | One-way ANOVA, post-hoc Dunnett's | Constant variance and approximate normality | Shapiro-Wilk (normality), Brown-Forsythe (eq. var.) |
| **SI 9** | 3 (1x macrophage stimulation) | 10 | 3 | 10 | One-way ANOVA, post-hoc Dunnett's | Constant variance and approximate normality | Shapiro-Wilk (normality), Brown-Forsythe (eq. var.) |
